# A Network of Noncoding Regulatory RNAs Acts in the Mammalian Brain

**DOI:** 10.1101/279687

**Authors:** Benjamin Kleaveland, Charlie Y. Shi, Joanna Stefano, David P. Bartel

## Abstract

Noncoding RNAs (ncRNAs) play increasingly appreciated gene-regulatory roles. Here, we describe a regulatory network centered on four ncRNAs—a long ncRNA, a circular RNA, and two microRNAs—using gene editing in mice to probe the molecular consequences of disrupting key components of this network. The long ncRNA Cyrano uses an extensively paired site to miR-7 to trigger destruction of this microRNA. Cyrano-directed miR-7 degradation is much more efficient than previously described examples of target-directed microRNA degradation, which come from studies of artificial and viral RNAs. By reducing miR-7 levels, Cyrano prevents repression of miR-7–targeted mRNAs and enables the accumulation of Cdr1as, a circular RNA known to regulate neuronal activity. Without Cyrano, excess miR-7 causes cytoplasmic destruction of Cdr1as, in part through enhanced slicing of Cdr1as by a second miRNA, miR-671. Thus, several types of ncRNAs can collaborate to establish a sophisticated regulatory network.

**HIGHLIGHTS:** A long noncoding RNA, a circular RNA, and two microRNAs form a regulatory network

The Cyrano long noncoding RNA directs potent, multiple-turnover destruction of miR-7

Unchecked miR-7 prevents accumulation of Cdr1as circular RNA in cytoplasm of neurons miR-7 prevents this accumulation by enhancing the miR-671-directed slicing of Cdr1as

## INTRODUCTION

MicroRNAs (miRNAs) are abundant ~22-nt RNAs that are required for normal mammalian development and dysregulated in human diseases such as cancer (Olive et al., 2015; Vidigal and Ventura, 2015). miRNAs regulate gene expression by base-pairing to mRNAs to induce mRNA decay and inhibit translation (Jonas and Izaurralde, 2015). Most miRNAs are extremely stable, with half-lives of days to weeks (Bail et al., 2010; Duffy et al., 2015; Gantier et al., 2011; Guo et al., 2015), which is attributed to their organization within the effector protein Argonaute. In Argonaute, the miRNA 5′ and 3′ ends are embedded in respective pockets that protect them from exonucleases that efficiently degrade naked single-stranded RNAs in the cell (Elkayam et al., 2012; Nakanishi et al., 2012; Schirle and MacRae, 2012). Nonetheless, some miRNAs are destabilized, either constitutively or in response to either neuronal excitation or growth factors (Avraham et al., 2010; Hwang et al., 2007; Krol et al., 2010; Rissland et al., 2011).

One way that miRNAs are destabilized is through the phenomenon of target RNA– directed miRNA degradation (TDMD). Artificial targets with extensive complementarity to the miRNA can trigger TDMD through a process that is associated with tailing (i.e., the addition of untemplated nucleotides) and trimming of the miRNA 3′ terminus but is otherwise poorly understood (Ameres et al., 2010; Baccarini et al., 2011; de la Mata et al., 2015; Denzler et al., 2016; Xie et al., 2012). Murine cytomegalovirus and herpesvirus saimiri (endemic in squirrel monkeys) have independently evolved RNAs that use this mechanism to promote the degradation of a host miRNA, miR-27, and a noncoding region within an mRNA of human cytomegalovirus induces TDMD of two related host miRNAs, miR-17 and miR-20a (Cazalla and Steitz, 2010; Libri et al., 2012; Marcinowski et al., 2012; Lee et al., 2013). In addition, a cellular transcript with TDMD activity has recently been reported; this transcript limits miR-29b expression in cerebellar granule neurons and regulates behavior in mice and fish (Bitetti et al., 2018).

Long noncoding RNAs (lncRNAs) are transcribed and processed like mRNAs but do not code for functional proteins. RNA sequencing has uncovered many thousands of lncRNAs, but the vast majority of these RNAs do not have known functions (Guttman and Rinn, 2012; Ulitsky and Bartel, 2013). The lncRNA Cyrano is particularly intriguing because it is broadly conserved in vertebrates and contains within its terminal exon a site with unusually high complementarity to the miR-7 miRNA (Ulitsky et al., 2011). Another class of abundant yet enigmatic RNAs is the circular RNAs (circRNAs), which are formed through the back-splicing of mRNA or lncRNA exons (Ebbesen et al., 2017). Among the most extensively characterized circRNAs is Cdr1as, a ~3 kb neuronally expressed circRNA. Cdr1as is conserved among mammals and dampens neuronal activity in mice (Piwecka et al., 2017). Interestingly, Cdr1as has many sites to miR-7 (130 and 73 sites in mouse and human Cdr1as, respectively), the same miRNA that pairs to Cyrano, and in vivo crosslinking data indicate that Cdr1as and Cyrano are the two RNAs most highly bound by miR-7 in human and mouse brain (Piwecka et al., 2017). Here, we show that Cyrano promotes the unusually efficient destruction of mature miR-7, which enables Cdr1as to accumulate in the mammalian brain.

## RESULTS

### Cyrano is a cytoplasmic lncRNA that promotes miR-7 destruction

In zebrafish, Cyrano is expressed throughout the developing nervous system (Ulitsky et al., 2011). To characterize Cyrano expression in mammals, we mined in situ hybridization and RNA-seq databases and performed RT–qPCR on 23 adult mouse tissues. In the mouse embryo, Cyrano was enriched in the early central and peripheral nervous systems (Figure S1A). In adult mice, Cyrano was ubiquitously expressed with 5–10-fold higher expression in brain regions and muscle (Figure 1A), a pattern that was largely conserved in human tissues (Figure S1B). Within the brain, Cyrano appeared more highly expressed in neurons than in glia (Figure S1A, S1C).

**Figure 1.**
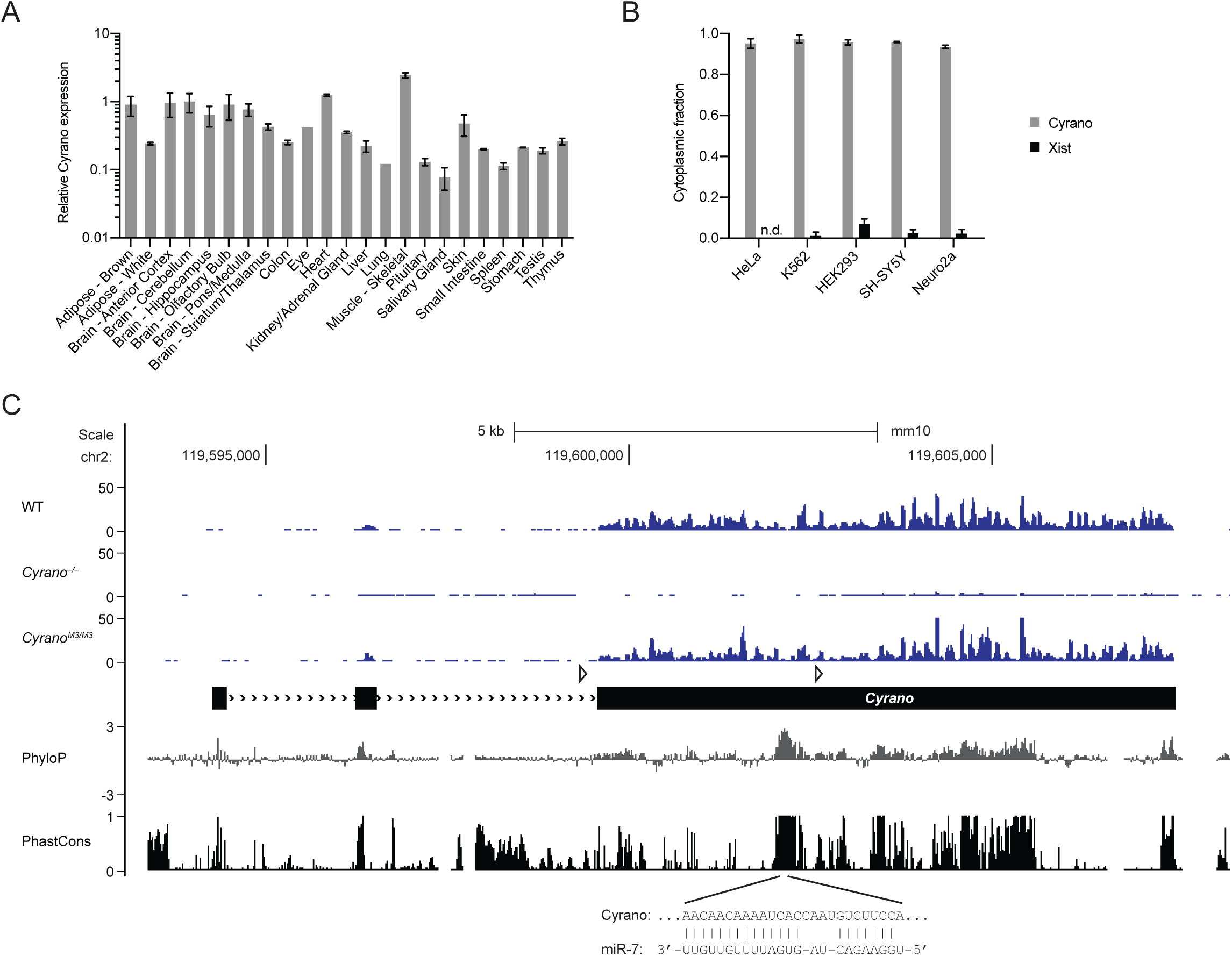
Characterization of Cyrano Expression. (A) Relative Cyrano expression in 23 tissues of 2-month-old mice. Shown are mean Cyrano levels, as determined by RT-qPCR, normalized to the geometric mean of the values for four housekeeping genes and relative to cerebellum (error bars, range; n = 1 biological replicate for eye and lung, n = 2 for the other 21 tissues). (B) Cyrano enrichment in the cytoplasm of human and mouse cell lines. Ratios were determined by RT-qPCR, using RNA from cytoplasmic and nuclear fractions, normalizing to a luciferase spike-in transcript (error bars, range; n = 2 biological replicates for HEK293, HeLa, and SH-SY5Y, n = 3 for K562, and Neuro2a). For comparison, results for the nuclear-localized Xist are also shown (n.d., not detected above background in cytoplasm). (C) Organization, conservation and expression of the murine Cyrano locus. Shown below the gene model (black boxes, exons; >>>, introns; open triangles, loxP sites), are conservation plots (PhyloP and PhastCons), which are based on a 40-genome placental mammal alignment generated relative to the mouse locus (Pollard et al., 2010; Siepel et al., 2005). Diagrammed below the conservation plots is the pairing of miR-7 to its highly complementary binding site, located within the most conserved region of Cyrano. Shown above the gene model are RNA-seq tracks for wild-type, *Cyrano*^*–/–*^ and *Cyrano*^*M3/M3*^ cerebellum (*y*-axis, reads) derived from libraries with similar sequencing depth.

Many well-characterized lncRNAs are found predominantly in the nucleus, playing roles in transcription, chromatin remodeling, splicing, rRNA processing, and nuclear organization (Guttman and Rinn, 2012; Engreitz et al., 2016). To assess the subcellular localization of Cyrano, we performed cytoplasmic–nuclear fractionation followed by RT–qPCR on several human and murine cell lines. In contrast to the nuclear lncRNA Xist, Cyrano was > 90% cytoplasmic in each cell line tested (Figure 1B).

To investigate the function of Cyrano in mammals, we generated Cyrano-deficient mice. The first half of exon 3, which includes the most highly conserved region of *Cyrano*, was targeted with flanking loxP sites in embryonic stem cells (Figure 1C, S1D). Two conditional *Cyrano* lines were established, and germline deletion was achieved by crossing with CMV-driven Cre mice. RNA-seq from Cyrano-deficient mice confirmed complete loss of the targeted region as well as a > 90% decrease in flanking Cyrano RNA, presumably due to destabilization caused by removal of the final splice-acceptor site (Figure 1C).

On both mixed SV129; C57Bl/6J and >99.5% C57Bl/6J backgrounds, Cyrano-deficient mice were born at the expected Mendelian frequency and had no overt gross or histologic abnormalities (Table S1, S2, data not shown). Cyrano-deficient adult mice had similar weights and one-year survival compared to sex-matched littermate controls, and basic neurobehavioral testing was unremarkable (data not shown). Nonetheless, Cyrano-deficient mice had some striking molecular phenotypes.

The most conserved region of Cyrano contains an unusual binding site for miR-7 (Figure 1C). In addition to pairing perfectly to the miRNA seed (nucleotides 2–7), which is a feature of effective canonical miRNA target sites (Bartel, 2009), this binding site has extensive pairing to the remainder of the miRNA (Ulitsky et al., 2011), with only a short internal loop that disrupts pairing to nucleotides 9 and 10 of the miRNA and should prevent Argonaute2-catalyzed slicing of Cyrano (Elbashir et al., 2001; Holen et al., 2002). This miRNA site architecture involving broadly conserved Watson–Crick pairing to all miRNA nucleotides except 9 and 10 is extremely rare in vertebrates and other bilaterian animals.

To determine whether Cyrano deficiency affected accumulation of miR-7 or any other miRNAs, we performed small-RNA sequencing on both wild-type and Cyrano-deficient brain samples. miR-7 is expressed from three independent loci in mice and humans. In humans, all three loci express identical mature miRNAs, whereas in mice they express two variants, miR-7a and miR-7b, which differ by a single nucleotide at position 10, with either possibility (A or U, respectively) forming a mismatch with the corresponding position of Cyrano. miR-7a and miR-7b were increased 40- and 47-fold in the cerebellum and 6- and 7-fold in the hippocampus (Figure 2A, 2B). Only two other miRNAs, miR-671 and miR-409-3p, were differentially expressed in both cerebellum and hippocampus, and the fold changes for these miRNAs were 1.4–2.8-fold.

**Figure 2.**
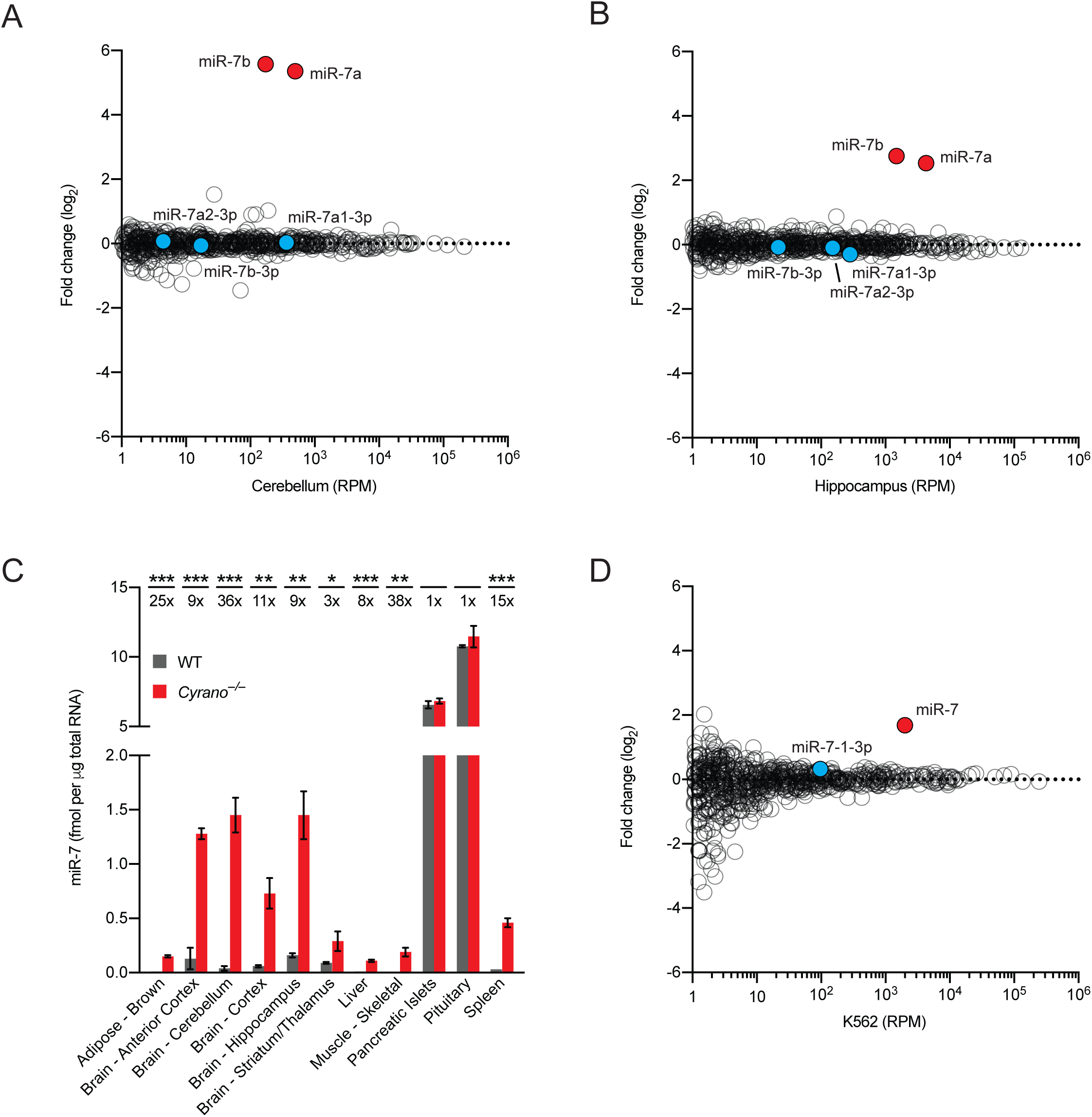
Cyrano Promotes miR-7 Destruction. (A) The influence of Cyrano on miRNA levels in the cerebellum. Shown are log_2_-fold changes in mean miRNA levels observed between wild-type and *Cyrano*^*–/–*^ cerebellum, as determined by small-RNA-seq and plotted as a function of expression in wild-type cerebellum (n = 3 biological replicates per genotype). Each circle represents one miRNA, showing results for all miRNAs expressed above 1 RPM (read per million miRNA reads) in wild-type cerebellum. Circles representing miR-7 paralogs and miR-7* strands are colored red and blue, respectively. (B) The influence of Cyrano on miRNA levels in the hippocampus. Otherwise, this panel is as in (A). (C) The influence of Cyrano on miR-7 levels in 11 mouse tissues. Plotted are the mean levels of miR-7 per μg total RNA for wild-type (gray bars) and *Cyrano*^*–/–*^ (red bars) tissues, as determined by quantitative RNA blots. For each tissue of each genotype, 3–4 biological replicates were analyzed alongside known quantities of miR-7 synthetic standards (error bars, standard deviation). Statistically significant fold changes are indicated (*, p < .05; **, p < .01; ***, p < .001, unpaired two-tailed *t*-test). (D) The influence of Cyrano on miRNA levels in K562 cells. Otherwise, this panel is as in (A).

In the canonical biogenesis pathway, each miRNA is transcribed by RNA polymerase II as a larger primary transcript with a distinctive hairpin structure that is cropped in the nucleus to generate a pre-miRNA hairpin and then further processed in the cytoplasm to generate the miRNA duplex (Ha and Kim, 2014). One strand of the duplex is preferentially loaded into Argonaute to form the silencing complex, whereas the passenger strand, referred to as the miRNA*, is degraded. In principle, increased miR-7 could be caused by 1) increased transcription of more than one miR-7 gene, 2) enhanced processing of the primary or precursor miR-7 RNAs, which is known to be highly regulated (Choudhury et al., 2013; Kumar et al., 2017; Lebedeva et al., 2011; Treiber et al., 2017), and/or 3) decreased degradation of miR-7a and miR-7b. To distinguish between these possibilities, we examined the levels of the intermediates in the miRNA-processing pathway. In contrast to the expectations for possibilities 1 and 2, we observed no changes in either the primary or precursor miR-7 RNAs in *Cyrano*^−/–^ mice (Figure S2A, data not shown). Most importantly, whereas possibilities 1 and 2 would each generate more of the miRNA duplex, our small-RNA sequencing results revealed no increase in the miRNA* strands of the miR-7 duplexes (Figure 2A, 2B, species with 3p suffixes), which strongly implied that Cyrano promotes degradation of the mature miR-7 after miR-7 is loaded into Argonaute.

To further characterize miR-7 expression in Cyrano-deficient mice, we performed quantitative RNA-blot analyses on samples from 11 adult tissues spanning the range of Cyrano expression (Figure 2C). In Cyrano-deficient pituitary and pancreatic islets, two tissues with relatively low expression of Cyrano and the highest baseline levels of miR-7, the miR-7 fold changes did not consistently exceed the variability observed among wild-type animals. Nonetheless, in the other Cyrano-deficient tissues tested, miR-7 levels significantly increased, with magnitudes ranging from 3–38 fold and increases most prominent in regions of Cyrano-deficient brain. Overall, miR-7 fold changes correlated with wild-type Cyrano expression across adult tissues (Figure S2B), as expected if Cyrano exerts greater effects in the tissues in which it is more highly expressed.

To determine when Cyrano begins to regulate miR-7, we examined the developing mouse brain beginning at E11.5. Increased levels of miR-7 were reliably observed as early as E13.5, although the fold changes (2–3 fold) were not as striking as those observed in most regions of the adult brain (Figure S2C). The ability of Cyrano to regulate miR-7 was not limited to the in vivo setting; miR-7 also increased 16–45 fold in primary cultures of mouse neurons derived from Cyrano-deficient animals (Figure S2D).

To test whether this regulation of miR-7 was conserved in humans, we used CRISPRi (Gilbert et al., 2013) to inhibit transcription from the *CYRANO* locus of human cells (also annotated as *OIP5-AS1*). CYRANO-depleted K562 cells showed a significant and specific ~3-fold increase in miR-7, with no significant change in miR-7-3p (Figure 2D). Similar results were also observed in additional human and mouse cell lines, with the increase in miR-7 correlating with the degree of CYRANO knockdown (Figure S2E). These findings confirmed that the ability of Cyrano to destabilize miR-7 is conserved in humans and requires transcription of the *CYRANO* locus.

### The miR-7 site in Cyrano directs multiple-turnover miR-7 destruction

Based on previous studies of TDMD triggered by artificial and viral transcripts, we suspected that the miR-7 binding site within Cyrano was crucial for the ability of this endogenous transcript to direct miRNA degradation. To test this, we used Cas9 to generate mice with small insertions and deletions in the Cyrano miR-7 site. In total, we isolated six strains, each homozygous for a different mutant allele (M1–M6), which were each deletions, except for one, which was an insertion/deletion (Figure 3A). Although some of the deletions were expected to disrupt miR-7 binding (Bartel, 2009), none affected expression of Cyrano in cerebellum and pituitary (Figure S3A–C), indicating that either miR-7 does not influence Cyrano accumulation or abrogated repression at this extensively paired site is offset by enhanced repression (due to increased levels of miR-7) at a second, more conventional seed site located at the beginning of exon 3. In contrast, all six mutations disrupted the ability of Cyrano to induce miR-7 degradation (Figure 3B). Three of the mutations (starred in Figure 3A) severely perturbed seed pairing with miR-7, and mice harboring these mutations showed increases in miR-7 levels resembling those of Cyrano-deficient mice, indicating that, as expected, seed pairing is critical for this interaction. More surprisingly, the M1 allele, which had the smallest perturbation, a 2-nt deletion eliminating only a single Watson-Crick pair to the seed region and reducing the size of the internal loop from 4 to 3 nucleotides, was also largely unable to promote miR-7 destruction. Overall, our findings showed that Cyrano-induced miR-7 degradation is exquisitely sensitive to the amount of Watson-Crick pairing (at least in the seed region) between Cyrano and miR-7. The importance of the miR-7 site also demonstrated that Cyrano acts through a form of TDMD reminiscent of that previously observed for artificial or viral transcripts.

**Figure 3.**
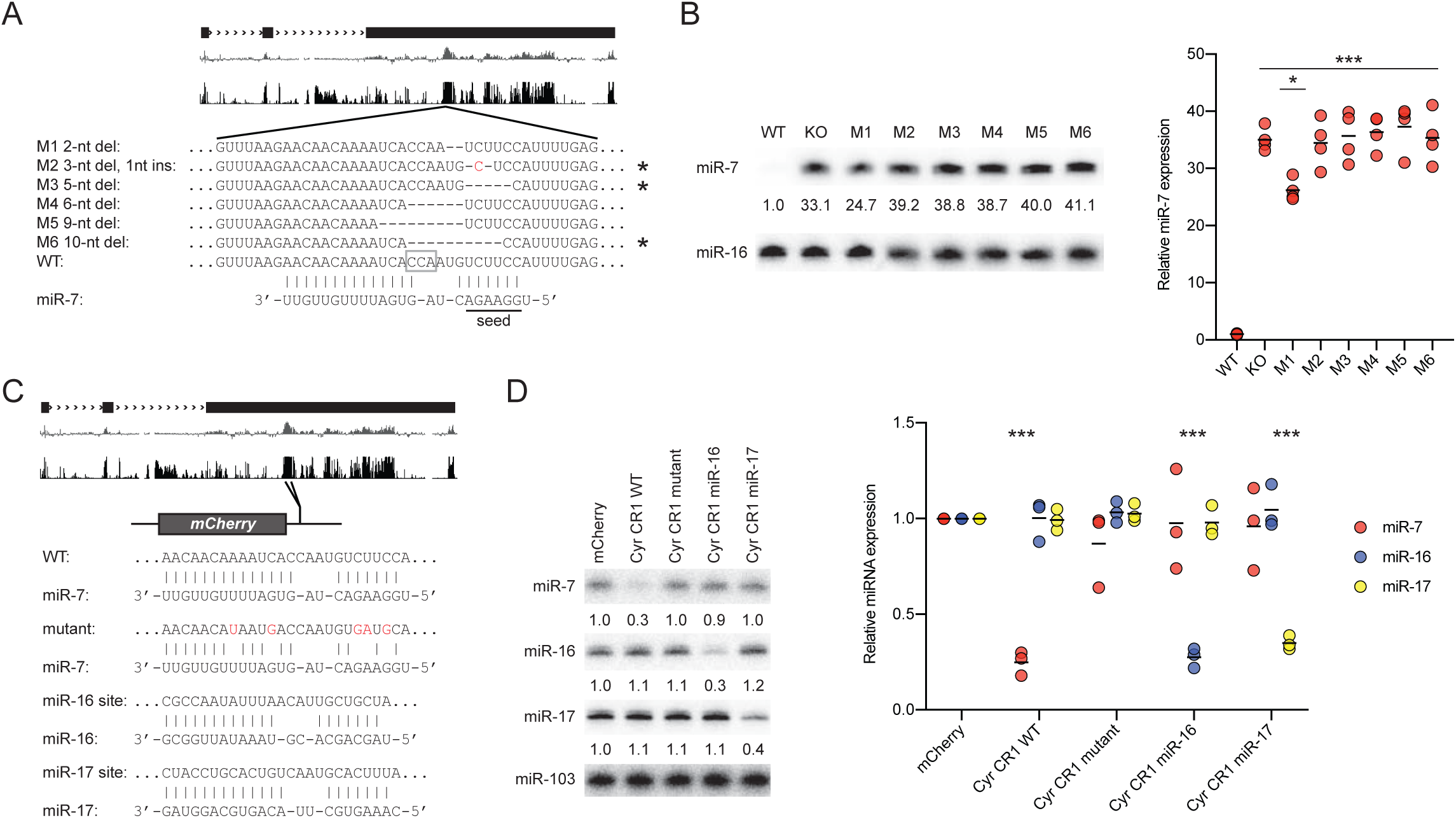
The miR-7 Site in Cyrano Specifies Destruction of miR-7. (A) Schematic of wild-type and Cyrano mutant alleles that change the extensively paired miR-7 site. Shown below the gene model and conservation tracks are the wild-type Cyrano miR-7 site and sites of six mutant alleles, M1–M6. The PAM motif used for Cas9-mediated mutagenesis is outlined (open gray box). Deleted nucleotides are indicated by dashes, and an inserted nucleotide is colored red. Alleles expected to disrupt pairing to the miR-7 seed are starred. (B) The importance of the miR-7 site for Cyrano function in cerebellum. At the left is a representative RNA blot measuring miR-7 levels in cerebellum of the indicated strains. In each lane, the level of miR-7 was normalized to that of miR-16 and reported relative to wild-type. Plotted at the right are relative miR-7 levels in the indicated strains (black bar, mean; n = 4 biological replicates). Statistically significant changes compared to wild type are indicated (***, p< 001, Tukey’s multiple-comparison test). The miR-7 level in M1 cerebellum also significantly differed from that in *Cyrano*^*–/–*^ and M2–M6 cerebellum (*, p < .05, Tukey’s multiple-comparison test). (C) Schematic of the wild-type CR1 (Cyrano conserved region) construct and engineered constructs that changed the miR-7 site. The miR-7 site in the CR1 was mutated to either disrupt pairing to miR-7 (substitutions indicated in red) or introduce a highly complementary site to either miR-16 or miR-17. (D) The importance of the complementary site for specifying CR1 function. At the left is a representative RNA blot measuring miR-7, miR-16, and miR-17 levels in HEK293T cells transfected with the constructs described in (C). Below each panel, the levels of the miRNA are reported relative to levels in the control mCherry transfection after normalizing to the level of miR-103. Plotted at the right are the relative levels of miR-7 (red circles), miR-16 (blue circles), and miR-17 (yellow circles) (black bars, mean; n = 3 biological replicates). Statistically significant changes compared to the mCherry control are indicated (***, p < .001, Dunnett’s multiple comparison test).

One important difference between the TDMD mediated by Cyrano and that described previously was the efficiency of the Cyrano-mediated phenomenon. In wild-type cerebellar granule neuron (CGN) cultures, Cyrano was present at 102 ± 16 molecules per cell, and miR-7 was present at 40 ± 7 molecules per cell (Table S3), whereas in Cyrano-deficient CGN, miR-7 levels increased 45 fold (Figure S2D) to 1800 molecules per cell. These values implied that Cyrano promoted TDMD with multiple turnover, with each Cyrano molecule responsible for the destruction of an average of ~17.6 miR-7 molecules. In contrast, similar calculations using values reported for previous examples of TDMD estimated the efficiency of the viral HSUR1 RNA to be ~0.1 miR-27 molecules per target (Cazalla and Steitz, 2010; Lee et al., 1988) and that of overexpressed synthetic single-site transcripts to be either 0.1–0.8 miR-122 molecules per target (Denzler et al., 2016), or 0.3–1.0 endogenous miRNA molecules per target, with multiple turnover achieved only when artificially boosting the miRNA level (de la Mata et al., 2015). These estimates indicated that Cyrano-promoted TDMD was 18–180-times more efficient than that of previously described examples.

To investigate further the specificity of this phenomenon, we examined whether the miR-7–destabilizing activity of Cyrano could be transferred to other miRNAs. For this, the 3′ UTR of an mCherry reporter gene was modified to include a 197-nt fragment from the most highly conserved region of Cyrano, using versions of this region that contained either the wild-type miR-7 site, a mutant miR-7 site, or an extensively complementary site to either miR-16 or miR-17 (Figure 3C). Decreased miR-7 was observed only in cells expressing the wild-type Cyrano region, whereas decreased miR-16 or miR-17 was observed specifically in cells expressing the region with cognate sites (Figure 3D). These results resembled those of analogous experiments that start with viral or artificial TDMD-triggering transcripts, and like those previous experiments, the amount of degradation was modest, with only ~3-fold reductions of the cognate miRNA observed for our constructs in HEK293T—much less than the 45-fold reduction observed for endogenous Cyrano in CGN culture (Figure S2D). Presumably, the neuronal setting (de la Mata et al., 2015). and other regions of the 8 kb Cyrano transcript were responsible for the unusually high efficiency of TDMD triggered by Cyrano in its endogenous context.

### Cyrano induces tailing and trimming of miR-7

Tailing and trimming of the miRNA 3′ terminus is often associated with TDMD (Ameres et al., 2010; Baccarini et al., 2011; Marcinowski et al., 2012; Xie et al., 2012; de la Mata et al., 2015; Denzler et al., 2016). To examine whether tailing and trimming was also associated with Cyrano-directed miR-7 degradation, we re-analyzed our small-RNA sequencing results, plotting the fraction of miR-7 reads of each length ranging from 19–30 nt. In both wild-type and Cyrano-deficient cerebellum, miR-7 predominantly existed as mature 23- and 24-nt isoforms (91–95% of all reads) (Figure 4A). However, a greater fraction of trimmed (≤ 22 nt) and tailed (≥ 25 nt) miR-7 reads were observed in the wild-type than in the Cyrano-deficient cerebellum. A similar pattern was observed in the hippocampus (Figure 4B). The trimmed miR-7 isoforms largely matched the genome-encoded miR-7 sequence, implying that tailing of trimmed species was either rare or destabilizing. When considering all trimmed fractions together, Cyrano increased the aggregate trimmed fraction 1.6 fold in cerebellum and 3.2 fold in hippocampus (Figure S4A, S4C). Similarly, Cyrano increased the aggregate tailed fraction 2.2 fold in cerebellum and 1.5 fold in hippocampus. Importantly, consistent differences in trimmed and tailed fractions were not observed for control miRNAs with comparable levels of expression, as exemplified by results for miR-92b (Figure 4A, 4B, S4A, S4C), and the same was true when analyzing all non-miR-7 reads in aggregate (Figure S4B, S4D).

**Figure 4.**
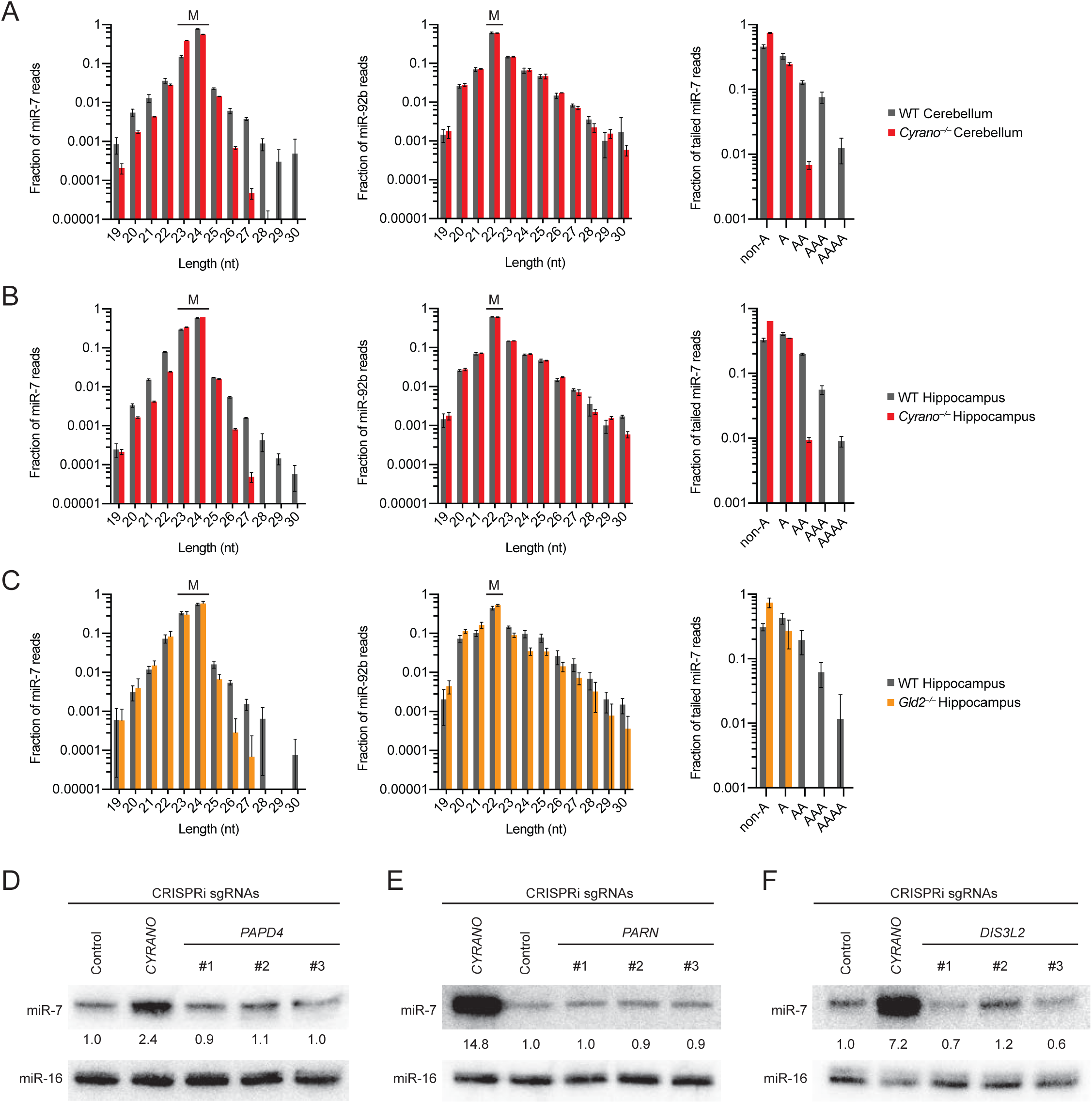
Tailing Can Be Uncoupled from miR-7 Trimming and Degradation. (A) The influence of Cyrano on miR-7 tailing and trimming in cerebellum. Plotted are length distributions of miR-7 reads (left) and miR-96b reads (middle) from wild-type (gray bars) and (B) *Cyrano*^*–/–*^ (red bars) cerebellum. The mature species for each miRNA are indicated (M). Also plotted are the fractions of tailed miR-7 reads that are mono- or oligoadenylylated (right) in wild-type (gray bars) and *Cyrano*^*–/–*^ (red bars) cerebellum. (C) The influence of Cyrano on miR-7 tailing and trimming in hippocampus. Otherwise, this panel is as in (A). (D) The influence of Gld2 on miR-7 tailing and trimming in hippocampus. Primary data from Mansur et al. (2016) were analyzed as in (A), with results for *Gld2^−/–^* hippocampus plotted with orange bars. Otherwise, this panel is as in (A). (E) The influence of PAPD4 on miR-7 expression in K562 cells. Shown are miR-7 levels, as determined by RNA blot, for stable cell lines expressing either a control sgRNA or an sgRNA targeting either *CYRANO* or *PAPD4* for CRISPRi-mediated knockdown. The levels of miR-7 are reported relative to the miR-7 level in the control cell line after normalizing to the level of miR-16. (F) The influence of PARN on miR-7 expression in K562 cells. Otherwise, this panel is as in (D). (G) The influence of DIS3L2 on miR-7 expression in K562 cells. Otherwise, this panel is as in (D).

In flies, tailing is primarily through addition of untemplated uridines (Ameres et al., 2010; Ameres et al., 2011), whereas in mammals, adenylylation appears more frequently than uridylylation, although both have been reported (Ameres et al., 2010; Baccarini et al., 2011; Marcinowski et al., 2012; Xie et al., 2012; de la Mata et al., 2015). Our analysis showed that Cyrano mostly increased the fraction of mature miR-7 reads with two or more untemplated adenosines, whereas monoadenylylation and tailing with other nucleotides were less affected (Figure 4A, 4B).

### Uncoupling tailing from trimming does not block degradation

Untemplated addition of nucleotides to the 3′ end of miR-7 could serve as a signal for an exonuclease to initiate 3′-to-5′ degradation, much like the addition of uridines to the 3′ end of pre-let-7 stimulates its destruction by Dis3l2 (Chang et al., 2013). Precedent for this idea also comes from observations of miR-21 and miR-122 in human leukemia and liver cell lines, respectively, where oligoadenylylated miR-21 and miR-122 isoforms are cleared by the cytoplasmic deadenylase Parn (Boele et al., 2014; Katoh et al., 2015). A prediction from this model was that disrupting the proteins that either add untemplated nucleotides or degrade miRNAs tailed with these nucleotides would disrupt Cyrano-directed degradation of miR-7.

Because Cyrano increased oligoadenylylation of miR-7, we hypothesized that the cytoplasmic polyA polymerase Gld2 might be responsible for this activity. Indeed, re-analyses of small-RNA sequencing data from wild-type and *Gld2^−/–^* hippocampus (Mansur et al., 2016) revealed that *Gld2* deletion resulted in a much lower proportion of tailed miR-7 reads and the disappearance of oligoadenylylated miR-7 reads (Figure 4C). These results for the *Gld2* deletion mirrored those observed for the *Cyrano* deletion, except the influence of Gld2 was not specific to miR-7, in that *Gld2* deletion decreased the proportion of tailed miRNA reads observed for a dozen miRNAs, including miR-92b (Figure 4C, S4E, S4F). Because *Gld2* deletion eliminated virtually all oligoadenylylation of miR-7, Gld2 must be responsible for the Cyrano-dependent oligoadenylylation of miR-7. However, unlike the *Cyrano*^*–/–*^ hippocampus, *Gld2^−/–^* hippocampus had no decrease in the proportion of trimmed miR-7 reads (Figure 4C, S4E). Moreover, loss of Gld2 did not significantly affect levels of any miRNA, including miR-7 (Mansur et al., 2016)(Figure S4G). Likewise CRISPRi stable knockdown of *PAPD4* (the human *Gld2* homolog) by as much as 98% in K562 cells did not affect miR-7 levels (Figure 4D, S4H).These results show that tailing (and more precisely, oligoadenylylation) induced by Cyrano can be uncoupled from both the trimming and the degradation of miR-7.

One way to reconcile the prevailing idea that tailing is required for TDMD with our observation that miR-7 levels remained unchanged after loss of Gld2 would be to propose that Gld2 has opposing consequences that somehow offset each other. Indeed, for miR-122, monoadenylylation by Gld2 stabilizes the miRNA, whereas oligoadenylylation by Gld2 destabilizes it by making it a substrate for PARN-catalyzed exonucleolytic decay (Katoh et al., 2015; Katoh et al., 2009). Because monoadenylylation of miR-122 predominates, Gld2 deletion in mice leads to an overall decrease in miR-122 in the liver (Katoh et al., 2009). Nevertheless, knockdown of PARN results in increased miR-122, revealing the destabilizing role for oligoadenylylation in this context (Katoh et al., 2015). To test if a similar phenomenon might be occurring for miR-7, we used CRISPRi to knockdown *PARN* in K562 cells. Stable knockdown of *PARN* by 99% had no effect on miR-7 levels (Figure 4E, S4I). Likewise, CRISPRi knockdown of *DIS3L2* by 98% had no discernable effect on miR-7 levels (Figure 4F, S4J). Although DIS3L2 is thought to prefer oligouridylylated substrates, a recent biochemical effort to identify factors involved in viral TDMD enriched for Dis3l2 (Haas et al., 2016). Taken together with our Gld2 results, these results indicate that miR-7 tailing, while clearly induced by Cyrano, is not on-pathway for Cyrano-directed miR-7 degradation, which calls into question the prevailing idea that tailing is an obligate step of TDMD.

### Loss of Cyrano causes increased miR-7 target repression

MicroRNAs bind to sites within the 3′ UTRs of mRNAs to promote mRNA destabilization and to a lesser extent translational repression (Eichhorn et al., 2014). The increased miR-7 from very low levels to intermediate levels in many Cyrano-deficient tissues (Figure 2C) raised the question of whether miR-7 targets might be more repressed in these tissues with increased miR-7. To answer this question, we performed RNA-seq on wild-type and Cyrano-deficient tissues and compared fold changes of predicted miR-7 targets to fold changes of mRNAs lacking a miR-7 site in their 3′ UTR. The effects on all predicted targets were compared to the effects on both conserved predictions and the top ~80 predictions (Agarwal et al., 2015), with the expectation that signal for repression would be greater when considering subsets predicted with higher confidence (i.e., top targets > conserved targets > all targets).

To provide context for the results for Cyrano-deficient tissues, we first processed RNA-seq data comparing wild-type to *Mir7a2^−/–^* pituitary (Ahmed et al., 2017). miR-7 is the most abundant miRNA in the pituitary, and deletion of *Mir7a2* leads to a massive reduction in miR-7 levels, an increase in predicted miR-7 targets, and infertility (Ahmed et al., 2017). Accordingly, we observed significant de-repression of predicted miR-7 targets in *Mir7a2^−/–^* pituitary, and the amount of derepression increased with higher-confidence predictions (Figure 5A). When analyzing Cyrano-deficient tissues, strong evidence for increased miR-7 target repression was observed in three of the ten tissues tested—striatum/thalamus, skeletal muscle, and pituitary (Figure 5B, 5C). The Cyrano-deficient spleen had an unanticipated increase in miR-7 targets, although this effect did not persist when considering only top targets. In tissues with increased miR-7 target repression, the amount of repression was substantially less than the effect of deleting miR-7 in the pituitary (Figure 5C), which is consistent with our results showing that the absolute levels of miR-7 in Cyrano-deficient tissues were still at least 7-fold lower than that of wild-type pituitary (Figure 2C). Despite the modest effect sizes, these results showed that by limiting miR-7 accumulation, Cyrano prevents repression of miR-7 targets in some tissues.

**Figure 5.**
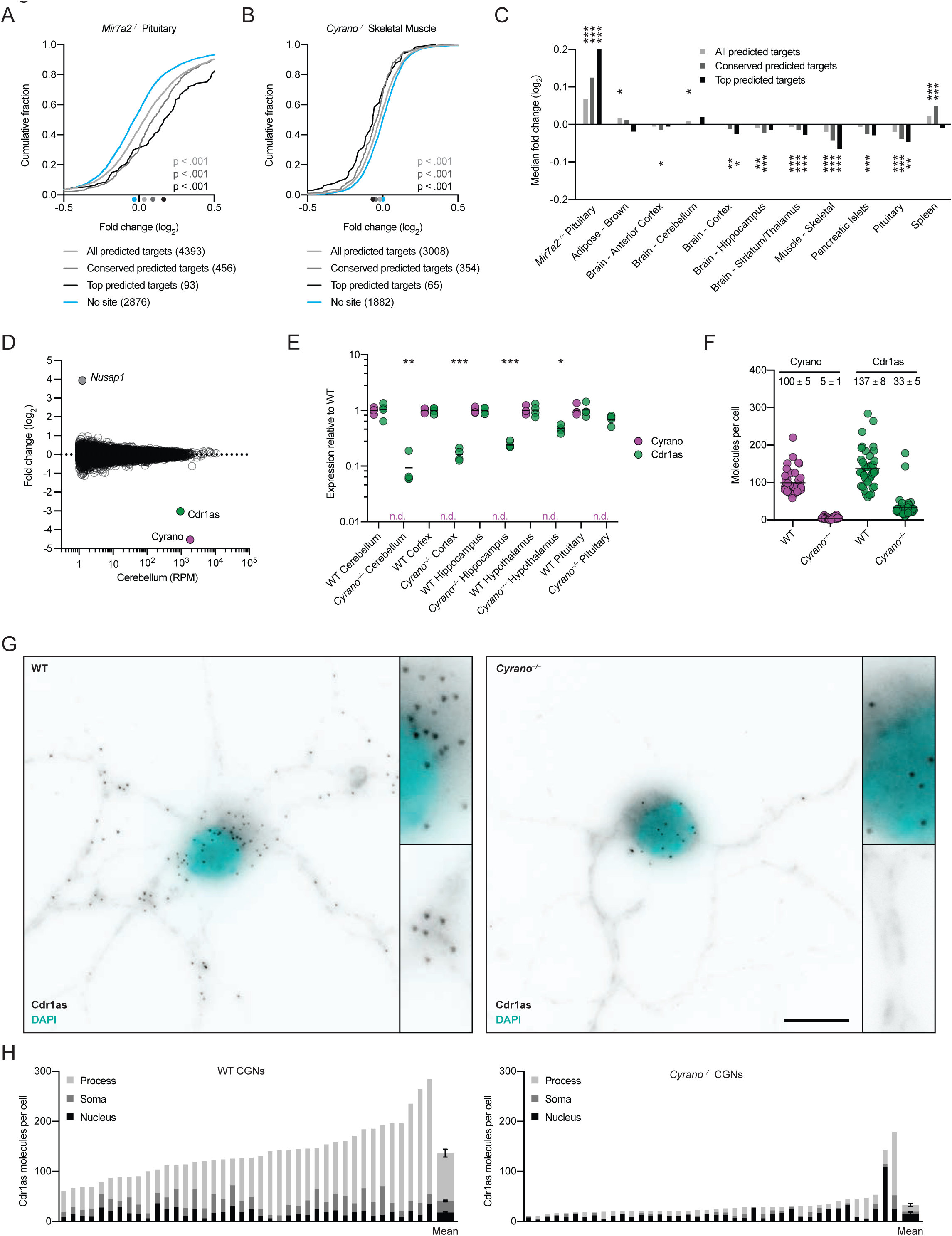
Cyrano Prevents Both the Repression of miR-7 Target mRNAs and the Destruction of Cdr1as. (A) The repression of predicted miR-7 targets by miR-7a-2 in the pituitary. Plotted are the cumulative distributions of mRNA changes observed after deleting *Mir7a2* in the pituitary (Ahmed et al., 2017), comparing the impact on all predicted miR-7 targets (light gray), conserved predicted miR-7 targets (dark gray), and top predicted miR-7 targets (black) to that of control mRNAs with no miR-7 site (blue). Median log_2_-fold changes for each set of mRNAs are indicated (colored dots below the *x*-axis). The statistical significance of differences between each set of predicted targets and the control mRNAs was determined by the Mann–Whitney test. (B) The influence of Cyrano on the predicted targets of miR-7 in skeletal muscle. Plotted are the cumulative distributions of mRNA changes observed in skeletal muscle after disrupting *Cyrano*. Otherwise, this panel is as in (A) The influence of Cyrano on the predicted targets of miR-7 in ten tissues. Plotted are median fold changes observed upon *Cyrano* disruption for the indicated sets of predicted miR-7 targets, after subtracting the median fold change of control genes. For reference, the effects of deleting *Mir7a2* in the pituitary, as evaluated in (A), are also shown (left-most set of bars). Statistically (C) significant changes from the control sets are indicated (*, p < .05; **, p < .01; ***, p < .001, Mann–Whitney test). (D) The influence of Cyrano on the expression of mRNAs and Cdr1as in cerebellum. Shown are log_2_-fold changes in mean RNA levels observed between wild-type and *Cyrano*^*–/–*^ cerebellum, as determined by RNA-seq and plotted as a function of expression in wild-type cerebellum (n = 4 biological replicates per genotype). Each circle represents a unique mRNA or noncoding RNA, showing results for all RNAs expressed above 1 RPM in wild-type cerebellum. Circles representing Cyrano, Cdr1as, and *Nusap1* are colored and labeled. (E) The influence of Cyrano on Cdr1as expression in brain and pituitary. Levels of Cyrano (purple) and Cdr1as (green), as determined by RT-qPCR, were normalized to that of *Actb* and plotted relative to mean wild-type expression (black bars, mean; n = 4 biological replicates per genotype; n.d., not detected). Statistically significant changes in Cdr1as levels in *Cyrano*^*–/–*^ tissues compared to wild-type tissues are indicated (*, p < .05; **, p < .01; ***, p < .001, unpaired two-tailed t-test). (F) The influence of Cyrano on Cdr1as expression in DIV11 CGNs, as determined by single-molecule FISH. Plotted are the number of Cyrano and Cdr1as molecules per cell for wild-type and *Cyrano*^*–/–*^ CGNs. Each circle represents the mean molecules per cell from a 100x microscopy field, each field containing 1–3 cells (n = 40 fields per genotype; black bars, mean). The mean of all cells of each genotype (± standard error) is also reported. (G–H) The influence of Cyrano on the accumulation of Cdr1as in neuronal cell bodies and processes. Shown in (G) are representative images from single-molecule FISH experiments probing for Cdr1as in wild-type (left) and *Cyrano*^*–/–*^ (right) CGNs, with insets showing increased magnification of portions of the soma and processes. RNA FISH signal is in black, cell nuclei in cyan (scale bar, 10 μm). Shown in (H) is quantitation of single-molecule FISH results obtained when probing for Cdr1as in wild-type (left) and *Cyrano*^*–/–*^ (right) CGNs. Each bar reports the mean number of molecules per cell observed in a 100x field containing 1-3 cells (n = 40 fields per genotype), differentiating molecules located in the nucleus (black), soma (excluding the nucleus, dark gray), and process (light gray). At the far right are bars plotting the means ± standard error.

### Cyrano enables the Cdr1as circRNA to accumulate in the brain

The signal for increased miR-7 repression observed in some Cyrano-deficient tissues was attributable to many miR-7 targets being repressed—but each by only a small amount. Indeed, when searching for any genes that on their own were differentially expressed in Cyrano-deficient cerebellum, with an adjusted *p* value < 0.01, only three genes were identified (Figure 5D). One was *Cyrano*, as expected (Figure 1C), and another was the downstream gene *Nusap1*, which increased due to read-through transcription and aberrant splicing (Figure S5A). Interestingly, the only other differentially expressed gene passing our *p* value cutoff was that of Cdr1as, the circRNA with 130 sites to miR-7. It decreased ~10-fold in *Cyrano*^*–/–*^ cerebellum (Figure 5D).

Because our RNA-seq libraries were generated after polyA-selection, which should not efficiently capture circular RNAs, we examined the Cdr1as profile to see if it corresponded to the linear transcript rather than the circular product. Very few reads mapped beyond the portion of Cdr1as known to circularize, suggesting that the RNA-seq was in fact capturing the circular isoform (Figure S5B). Moreover, RT–qPCR analysis using primers that spanned the back-splice junction (and were thus specific to the circular isoform) found a similar 10-fold reduction in Cdr1as in Cyrano-deficient cerebellum (Figure 5E). In wild-type mice, Cdr1as is expressed in neurons throughout the brain with highest expression in the cerebellum (Hansen et al., 2013; Memczak et al., 2013; Piwecka et al., 2017); Figure S5C). To determine if this effect on Cdr1as extended beyond the cerebellum, we also measured Cdr1as in other regions of the brain as well as the pituitary. Cdr1as decreased throughout the Cyrano-deficient brain, with the greatest effect observed in the cerebellum, followed by cortex, hippocampus, and hypothalamus (Figure 5E), mirroring the rank order of miR-7 fold changes in these regions (Figure 2C). No significant change was observed for Cdr1as expression in the pituitary.

To determine the subcellular location of Cyrano activity, we performed single-molecule FISH (fluorescence in situ hybridization), probing for Cyrano and Cdr1as in cultured CGNs, the most abundant type of neuron in the cerebellum. An average of 100 molecules of Cyrano and 137 molecules of Cdr1as were observed in wild-type CGNs (Figure 5F), which agreed with our RT–qPCR estimates (Table S3). Cyrano was observed predominantly in the cytoplasm (73%) with molecules distributed in both soma and processes (Figure S5D–E). As previously described in cortical neurons (Piwecka et al., 2017), Cdr1as was also predominantly localized to the cytoplasm (86%), with most molecules found in the processes (Figure 5G left, 5H left). In Cyrano-deficient CGNs, some transcripts were detected, as expected for a probe set that extended beyond the deleted portion of Cyrano, and compared to wild type, the number of molecules detected was greatly reduced (Figure S5D–E), consistent with the RNA-seq results showing a large decrease in Cyrano RNA flanking the deletion (Figure 1C). The number of Cdr1as molecules also decreased in nearly all neurons (Figure 5F), and strikingly, this decrease in Cdr1as was entirely attributable to depletion from the soma and processes (Figure 5G right, 5H right). The preservation of Cdr1as levels in the nucleus suggested that the large reduction of Cdr1as in Cyrano-deficient cells occurred through cytoplasmic destruction of Cdr1as rather than reduced Cdr1as production or defective export. These results suggested a model in which high levels of miR-7 induce cytoplasmic destruction of Cdr1as, and Cyrano, by limiting miR-7, enables Cdr1as to accumulate in neuronal cell bodies and processes.

### Cdr1as is regulated by miR-7 and miR-671

To explore further the idea that miR-7 links the fate of Cyrano to that of Cdr1as, we analyzed RNA-seq results from the cerebellum of *Cyrano*^*M3/M3*^ mice, which were homozygous for a 5-nt deletion in the miR-7 complementary site (Figure 3A). When compared to results from wild-type cerebellum, the expression of *Cyrano* did not change (Figure 6A, 1C), as expected from our previous RT–qPCR analyses of these mice (Figure S3A–C). Expression of the downstream gene *Nusap1* also did not change, as expected in this mutant predicted to retain normal Cyrano splicing and 3′-end processing (Figure 6a). Remarkably, *Cdr1as* was the only differentially expressed gene identified with adjusted *p* value < 0.01, and its expression decreased ~10-fold in *Cyrano*^*M3/M3*^ cerebellum, which resembled the decrease observed in the *Cyrano*^−/–^ cerebellum. This decrease in Cdr1as was corroborated by RT–qPCR analyses of cerebellar samples from all six lines with Cyrano miR-7 site mutations (Figure S6A). These results showed that the miR-7 complementary site within Cyrano was required for this lncRNA to influence Cdr1as accumulation.

**Figure 6.**
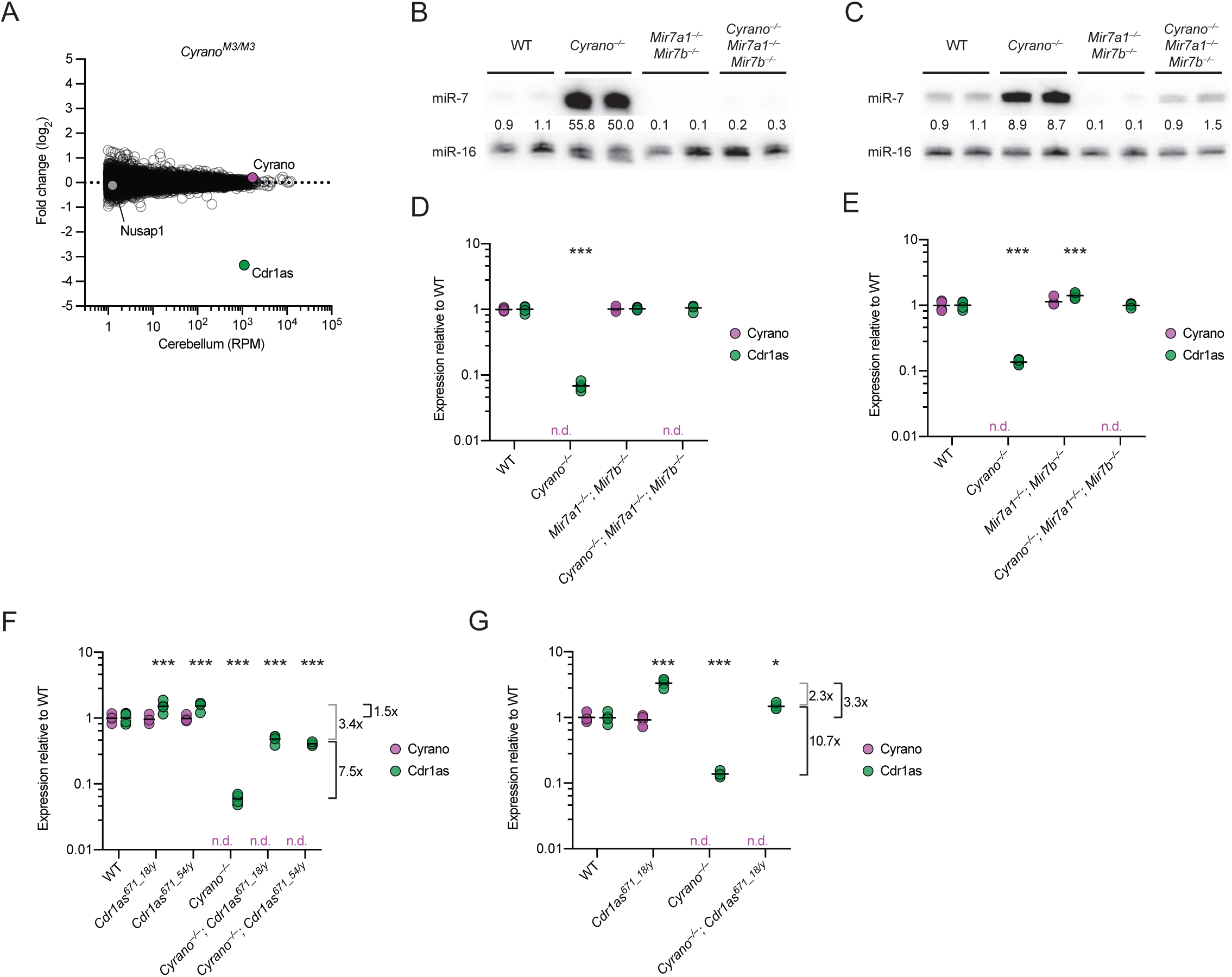
Cdr1as is Regulated by miR-7 and miR-671 (A) The importance of the miR-7 complementary site for the function of Cyrano in influencing Cdr1as levels in cerebellum. Shown are log_2_-fold changes in mean RNA levels between wild-type and *Cyrano*^*M3/M3*^ cerebellum, as determined by RNA-seq and plotted as a function of expression in wild-type cerebellum (n = 3–4 biological replicates per genotype). Otherwise, this panel is as in Figure 5D. (B–C) Substantially reduced miR-7 expression in cerebellum and cortex of *Mir7a1^−/–^*; *Mir7b^−/–^* mice, even without functional Cyrano. Shown are RNA blots measuring miR-7 levels in the cerebellum (B) and cortex (C) of mice with the indicated genotypes. Levels of miR-7 were normalized to those of miR-16 and are reported relative to the mean wild-type level. (D–E) A requirement of miR-7 for the Cdr1as reductions observed in mice without functional Cyrano. Cyrano (purple) and Cdr1as (green) levels in cerebellum (D) and cortex (E) of mice with the indicated genotypes were determined by RT-qPCR, normalized to *Actb*, and plotted relative to mean wild-type expression (black bars, mean; n = 4 biological replicates per genotype; n.d., not detected). Statistical significance of the differences in Cdr1as level in each mutant tissue compared to that in wild-type tissue is indicated (***, p < .001, Tukey’s multiple-comparison test).(F–G) The influence of the miR-671 site within Cdr1as on Cdr1as levels both with and without functional Cyrano. Cyrano and Cdr1as in cerebellum (D) and cortex (E) of mice with the indicated genotypes were measured and plotted as in (D–E). The mean fold changes in Cdr1as attributable to the miR-671 site, either with or without functional Cyrano (black brackets), and not attributable to the miR-671 site (gray brackets) are indicated to the right of each plot.

To test whether miR-7 is required for the destabilization of Cdr1as in the absence of Cyrano, we generated mice that lacked the two miR-7 paralogs most highly expressed in the brain (Figure S6B). These *Mir7a1^−/–^*; *Mir7b^−/–^* mice were born at the expected Mendelian frequency and had no gross abnormalities (Table S4). Although mice with cortex-specific overexpression of a miR-7 sponge develop smaller brains (Pollock et al., 2014), we did not observe any difference in brain size between wild-type, double-heterozygous, and double-knockout mice (Figure S6c). These mice did have reduced fertility, but as expected for deletion of these paralogs that were not as highly expressed in the pituitary, this defect was not a severe as that observed for *Mir7a2^−/–^* (Ahmed 2017), and most *Mir7a1^−/–^*; *Mir7b^−/–^* males and females were able to mate successfully.

Analysis of several brain regions using RNA blots showed that miR-7 levels substantially decreased throughout the brain of *Mir7a1^−/–^*; *Mir7b^−/–^* animals (Figure 6B, 6C, S6D). Global analyses of predicted miR-7 targets revealed no significant changes in cerebellum and cortex, which indicated that the absolute levels of miR-7 normally found in these tissues are too low to direct detectable target repression (Figure S6E). When combining the *Cyrano* deletion with the *Mir7* double knockout to generate triple-knockout mice (*Cyrano^−/–^; Mir7a1^−/–^; Mir7b^−/–^*), miR-7 levels remained at or below those of wild-type mice (Figure 6B, 6C), thus providing a context in which Cyrano loss is no longer tied to abnormally high miR-7 levels. In this context, Cdr1as was insensitive to Cyrano loss, demonstrating that increased miR-7 levels were required for Cdr1as destabilization in the absence of Cyrano (Figure 6D, 6E). Altogether, our results strongly support a model in which Cyrano-directed miR-7 degradation protects Cdr1as from miR-7– mediated degradation, thereby allowing Cdr1as to accumulate in neuronal cell bodies and processes.

In addition to many conserved miR-7 binding sites, Cdr1as also contains a single conserved site to miR-671, which pairs to the miRNA with complementarity sufficient to trigger Argonaute2-catalyzed slicing of the circRNA (Hansen et al., 2011). We hypothesized that miR-7–mediated degradation of Cdr1as might occur through enhanced miR-671 binding and/or cleavage of Cdr1as. To be able to test this hypothesis, we generated mice with mutations in the Cdr1as miR-671 site (Figure S6F) and selected for further study mutant alleles with either 18- or 54-nt deletions disrupting the miR-671 site. Hemizygous males (*Cdr1as* is on the X chromosome) and homozygous females with either of these deletions were born at the expected Mendelian frequencies, with no changes observed in miR-7 levels (Tables S5, S6; Figure S6G). Deletion of the miR-671 site within Cdr1as led to increased levels of Cdr1as in the brain, up to ~4-fold in the cortex and hippocampus (Figure S6H). These results indicate that miR-671–mediated slicing of Cdr1as, known to occur in cultured cells (Hansen et al., 2011), also occurs in the animal, and indeed is sufficiently robust to set the Cdr1as levels in different regions of the brain.

With these *Cdr1as^671^* strains in hand, we generated *Cyrano*^*–/–*^; *Cdr1as^671/y^* mice and, after confirming that the *Cyrano* deletion in the *Cdr1as^671^* background had increased miR-7 levels (Figure S6I), we assessed Cdr1as expression by RT–qPCR. A full rescue of Cdr1as expression in *Cyrano*^*–/–*^; *Cdr1as^671/y^* mice was expected to increase Cdr1as to the same level as that of *Cdr1as^671/y^* mice, whereas no rescue was expected to increase Cdr1as relative to the *Cyrano*^*–/–*^ by a fold change similar to that which was observed between wild-type and *Cdr1as^671/y^* mice (1.5 and 3.3 fold in cerebellum and cortex, respectively). Instead of either of these extremes, we observed that Cdr1as levels were partially restored. In cerebellum, for example, Cdr1as increased 7.5 fold, implying a 5 fold (7.5 / 1.5 fold) increase in slicing at the miR-671 site in the absence of Cyrano activity (Figure 6F). The observation that preventing miR-671–directed slicing only partially restored Cdr1as levels indicated that at least one other miR-7–dependent mechanism acts to degrade Cdr1as in this tissue, with a presumed 3.4-fold effect. Likewise, in cortex, Cdr1as increased 10.7 fold, implying a 3.2 fold (10.7 / 3.3 fold) increase in slicing at the miR-671 site, with another 2.3 fold attributed to another mechanism (Figure 6G). Taken together, our results show that in the absence of Cyrano-directed miR-7 degradation, miR-7 levels in brain tissues increase such that miR-7 destroys Cdr1as through at least two mechanisms, one of which increases slicing at the miR-671 site.

## DISCUSSION

We characterize a posttranscriptional regulatory network centered on four noncoding RNAs— one lncRNA, one circRNA, and two miRNAs—using a panel of mouse knockouts to disrupt nodes and edges in the network and thereby learn their molecular consequences in the animal (Figure 7). We find that the conserved lncRNA Cyrano functions as an endogenous and potent miRNA assassin. In targeting miR-7 for degradation, Cyrano prevents miR-7 from repressing its mRNA targets in some tissues, including skeletal muscle and striatum/thalamus. More strikingly, Cyrano also prevents miR-7 from causing the efficient cytoplasmic destruction of the circRNA Cdr1as in many brain tissues, which occurs in part by enhancing miR-671–directed slicing of Cdr1as.

**Figure 7.**
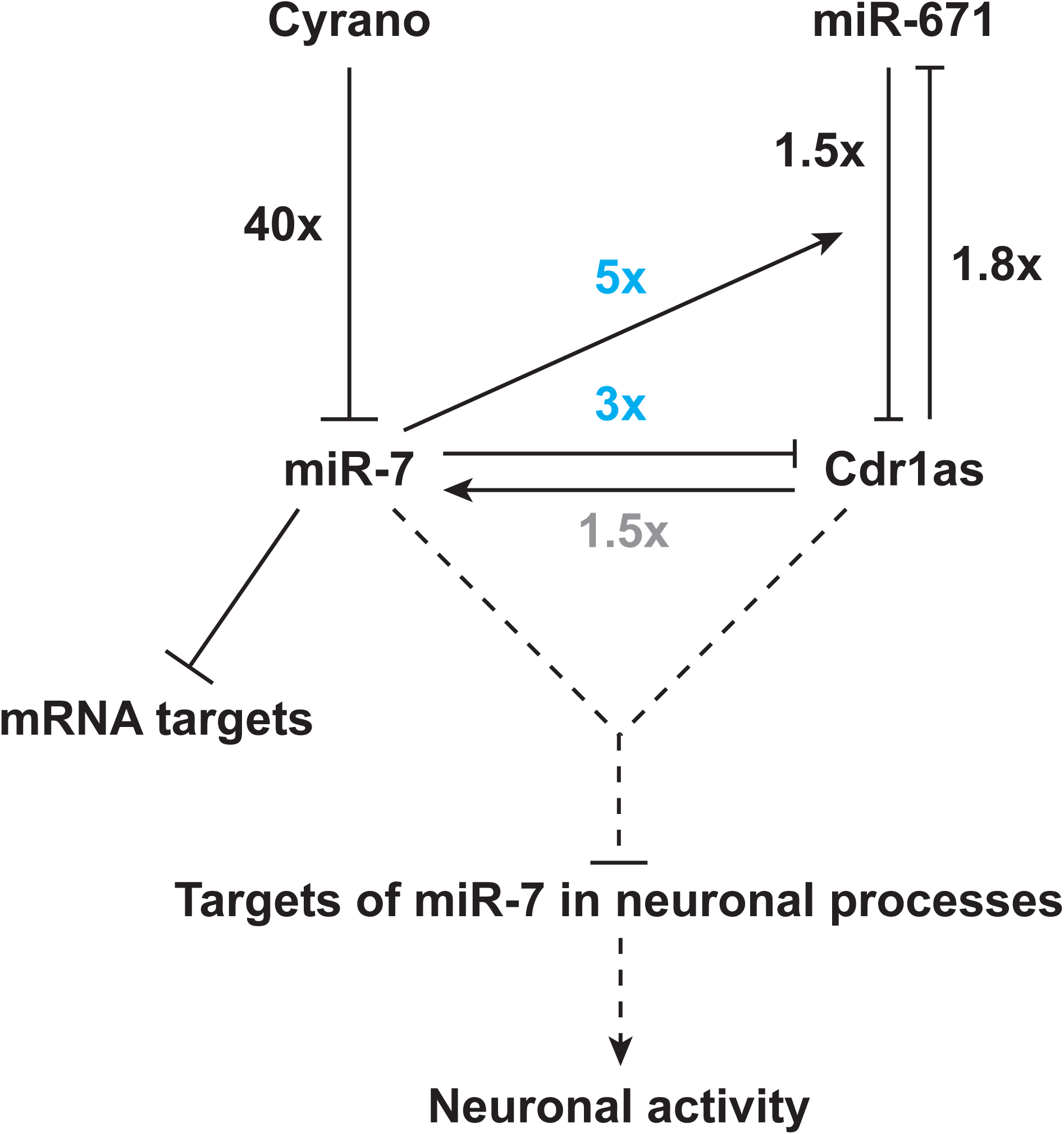
A Network of Noncoding Regulatory RNAs Acts in the Mammalian Brain For each interaction, the effect was measured using a mouse-knockout model that disrupted either the relevant ncRNA (either Cyrano, miR-7, or Cdr1as) or miRNA-binding site (either the miR-7 site in Cyrano or the miR-671 site in Cdr1as). The fold changes reported are from the cerebellum, observed either in this study (black or blue) or previously (gray, Piwecka et al., 2017). The 1.8-fold increase in miR-671 observed in our mice with the disrupted Cdr1as miR-671 site was similar to that observed in *Cdr1as*-knockout mice (Piwecka et al., 2017). Regulation that emerges with derepression of miR-7 (as opposed to deletion of miR-7) is indicated (light blue fold changes). Repression of miR-7 targets is also depicted, but without a fold change because it was detected only in other tissues.

### Insights into target RNA-directed miRNA degradation

One surprising aspect of Cyrano-targeted miR-7 destruction is its potency—occurring with multiple turnover, 18-180 fold more efficiently than previously described TDMD examples. Although a highly complementary miRNA binding site is required and often sufficient to induce miRNA degradation (Ameres et al., 2010), it is not the whole story. For example, analysis of the human CMV RNA shows that a 16-nt sequence downstream of the complementary site is important for recapitulating the miRNA degradation observed with the full-length RNA (Lee et al., 2013). The ~8 kb Cyrano transcript is substantially longer than viral transcripts that direct miRNA degradation and has deeply conserved blocks of sequence not only proximal to the miR-7–complementary site but also elsewhere within the transcript (Ulitsky et al., 2011)(Figure 1C). Some of these regions might be conserved because they enhance the efficiency of Cyrano-directed miR-7 degradation, presumably by helping recruit miR-7–programmed Argonaute or miRNA-degradation factors. Such a function for the more distal conserved regions would help explain why we observed only a 3-fold decrease in miR-7 after > 100-fold overexpression of the 197-nt site-containing region of Cyrano (Figure 3E, 3F). This strategy of using multiple regions of a long ncRNA to boost degradation efficacy would be suitable for an endogenous transcript, whereas the pressure to minimize genome size might favor the viral strategy of using a larger number of shorter, less efficient transcripts.

Another factor that presumably boosts the efficiency of Cyrano is cellular context. The most efficient TDMD previously observed was in neurons (de la Mata et al., 2015), and we observed the most potent Cyrano activity (98% miR-7 reduction, with ~17 molecules degraded per Cyrano molecule) in cultured neurons (Figure S2D). Moreover, our analyses of *Cyrano*^*–/–*^ tissues suggest that, in addition to neurons, skeletal muscle and brown fat are cell types that provide unusually favorable contexts for TDMD (Figure 2C). Perhaps these cell types express more of a factor that is limiting for efficient TDMD.

What then might be the elusive protein factors responsible for TDMD? As tailing and trimming of miRNA 3′ ends co-occur with TDMD, an attractive model has been that tailing of the 3′ end of the miRNA licenses the miRNA for degradation by a 3′-to-5′ exonuclease. This model is consistent with findings that 1) 2’-O-methylation, a modification that both prevents 3′ tailing and stabilizes small RNAs, is typically found on plant miRNAs and other classes of small regulatory RNAs that have extensive pairing to most targets, yet is absent on metazoan miRNAs, which do not have extensive pairing to most targets (Ameres et al., 2010; Ji and Chen, 2012), 2) tailing of certain metazoan miRNAs and pre-miRNAs promote their destruction (Boele et al., 2014; Chang et al., 2013; Katoh et al., 2015), and 3) nucleotidyltransferases and 3′-to-5′ exonucleases co-purify and cooperate in protein complexes (Kim and Richter, 2006; Reimao-Pinto et al., 2016). Despite the appeal of this model, our findings comparing *Cyrano*^*–/–*^ and *Gld2^−^/–* mice revealed that tailing can, in fact, be uncoupled from trimming and degradation. If tailing is off-pathway for degradation, trimming might also be an off-pathway event caused by transiently freeing the 3′ end of the miRNA from its protected pocket in Argonaute, in which case, TDMD need not occur through a 3′-to-5′ exonuclease. Indeed, we found that severe knockdown of candidate 3′-to-5′ exonucleases had no detectable effect on Cyrano-directed miR-7 degradation. Perhaps the key factors recruited by Cyrano are not nucleases at all but instead act on Argonaute to either degrade the protein or help eject the Cyrano-bound miR-7. Once removed from the protective grip of Argonaute, miR-7 could be destroyed redundantly by any number of cellular nucleases. The discovery of the unusually efficient TDMD mediated by Cyrano provides a promising new system for experimental dissection of this phenomenon.

### Destruction of a circRNA

Although much has been learned about circRNA biogenesis, less is known about how circRNAs are degraded and whether circRNA turnover is regulated (Ebbesen et al., 2017). CircRNAs are much more stable than mRNAs, presumably due to the absence of free 5′ and 3′ ends (Jeck et al., 2013; Memczak et al., 2013; Enuka et al., 2016). They are also expected to be resistant to miRNA-mediated RNA destabilization, which occurs through deadenylation and decapping (Jonas and Izaurralde, 2015). Cdr1as is unusual in that it has a built-in mechanism for destruction via miR-671–directed slicing (Hansen et al., 2011). We found that this mechanism is active in the animal, controlling the steady-state accumulation of Cdr1as in the brain, especially in the cortex. In *Cyrano*^*–/–*^ brains, miR-671–directed slicing of Cdr1as is further enhanced due to increased levels of miR-7. Perhaps miR-7, which has sites within close proximity of the miR-671 site, helps recruit or retain the miR-671 silencing complex, through an undefined mechanism that also explains the cooperative effects of closely spaced sites in mediating repression (Grimson et al., 2007; Saetrom et al., 2007). For example, the miR-7 and miR-671 silencing complexes might cooperatively bind to Cdr1as, or binding of miR-7 to Cdr1as might change the conformation of Cdr1as so as to increase accessibility of the miR-671 site.

Increased miR-671–directed slicing explains 3–5 fold of the Cdr1as degradation observed in *Cyrano*^*–/–*^ cerebellum and cortex; at least one other miR-7–dependent mechanism is responsible for the remaining 2–3 fold. miR-7 is not expected to direct slicing of Cdr1as by Argonaute2 because Cdr1as has no site with extensive complementarity to miR-7. Perhaps, Cdr1as studded with too many Argonautes is trafficked to a more degradative subcellular compartment or environment.

### A network of noncoding RNAs acting in neurons

What might be the purpose of this baroque posttranscriptional regulatory network centered on these four noncoding RNAs? With its unusually high number of miR-7 binding sites, Cdr1as was initially proposed to act as an inhibitor of miR-7, titrating miR-7 away from mRNA targets while resisting deadenylation and decapping typically induced by stable Argonaute binding (Hansen et al., 2013; Memczak et al., 2013). If Cdr1as does act as a miR-7 sponge, then deleting it should lead to decreased levels of miR-7 targets. However, such a decrease is not observed in the brains of *Cdr1as*-knockout animals (Piwecka et al., 2017). Instead, the role of Cdr1as in dampening neuronal activity in the brain is partly attributed to the idea that it protects miR-7 from degradation thereby causing increased repression of miR-7 targets (Piwecka et al., 2017).

One concern with this model is the low magnitude of the changes in conserved predicted miR-7 targets attributed to the = ≤ 2-fold reduction in miR-7 in *Cdr1as*-knockout tissues (Piwecka et al., 2017). The results of our experiments are also difficult to reconcile with the notion that the modest decrease in miR-7 (= ≤2-fold) observed in the *Cdr1as* knockout helps explain the knockout phenotype. When examining the effects of increased miR-7 after disrupting Cyrano, we did not find evidence for increased repression of miR-7 targets in regions of the brain where Cdr1as is thought to be active, such as cerebellum and cortex (Figure 5C). More importantly, when examining the effects of decreased miR-7 levels (> 5 fold) in the *Mir7* double knockout, we did not find evidence for decreased repression of miR-7 targets in those same brain regions (Figure S6I). Together with our miR-7 expression measurements, these results seem to indicate that in these tissues in which Cdr1as is exerting its influence on neuronal activity, miR-7 is expressed at levels too low to detectibly influence its set of predicted targets.

Nonetheless, we found no reason to return to the original idea that Cdr1as acts primarily as an inhibitor of miR-7. Indeed, we found the opposite—that miR-7 can act to inhibit Cdr1as.Moreover, our results showing the potent TDMD activity of Cyrano do support one aspect of the recently proposed model, which is the idea that Cdr1as might protect miR-7 from a competing degradative activity. For example, in a typical CGN cell, the 18,000 miR-7 sites of the ~140 Cdr1as molecules would add to the 50,000 miR-7 sites in the transcriptome (effective target-site concentration of the transcriptome, estimated as in Denzler et al., 2016) to increase competition with the ~100 Cyrano sites, thereby rescuing from Cyrano-directed degradation a slightly higher fraction of the ~1,800 miR-7 molecules made in these cells (Table S3). In this way, Cdr1as could be responsible for increasing the miR-7 molecules in the cell—from 25–30 molecules to the observed 40 (Table S3).

So how else might Cdr1as function to dampen neuronal activity? The many conserved sites to miR-7 suggest that Cdr1as activity somehow involves this miRNA. Although we do not favor the idea that its influence on miR-7 levels is at the heart of its function, both the rejection of its role as a miR-7 inhibitor and the presence of its activity in tissues in which miR-7 activity seems too low to have any consequence on its general target repertoire, do imply that Cdr1as might potentiate miR-7 activity in some other way. We favor the proposal that Cdr1as acts to engineer spatial and perhaps even temporal control over the delivery of miR-7 in neurons (Piwecka et al., 2017). Controlled repression of miR-7 targets at synapses (or a subset of synapses) might limit neuronal activity without detectable effects on the bulk of the miR-7 targets. This idea is consistent with the observation that miR-7 has multiple conserved targets involved in vesicle trafficking in the pancreas (Latreille et al., 2014) and also consistent with the enrichment of miR-7 in processes, as inferred from biochemical fractionation studies comparing synaptosomes with bulk brain tissue (Siegel et al., 2009; Smalheiser et al., 2014).

Another fascinating aspect of this network is its construction, built from components with varying depths of conservation. Among the four ncRNAs, miR-7 is the oldest, having descended from a common ancestor of all bilaterian animals, with its robust expression in neurosecretory brain centers conserved from annelids to humans (Tessmar-Raible et al., 2007). While invertebrates typically have a single copy of *Mir7*, vertebrates have 3–5 copies, located on different chromosomes (Fromm et al., 2015). Amplification of miR-7 coincided with the emergence of Cyrano, raising the possibility that Cyrano’s ancestral function might have been to minimize miR-7 activity in non-neuroendocrine tissues. It is within this context that both Cdr1as and miR-671 appeared in a common ancestor of placental mammals. That Cdr1as contains binding sites for miR-7 instead of another lowly expressed neuronal miRNA suggests that perhaps what matters is not merely the low expression of miR-7 but also its potent on–off switch in the form of Cyrano-mediated degradation. Knowledge of this network of noncoding RNAs, together with our knockout models disrupting nodes and edges of the network, help lay the foundation for experimental tests of Cdr1as molecular function and mechanism, which could ultimately provide a better understanding of the rationale for the emergence of this network.

## AUTHOR CONTRIBUTIONS

B.K. and D.P.B conceived the project and designed the study. B.K. performed all of the experiments, with contributions from J.S., except for the CRISPRi experiments, which were performed by C.Y.S. B.K. performed all of the analyses and drafted the manuscript. B.K., C.Y.S., and D.P.B revised the manuscript.

## ACKNOWLEDGMENTS

We thank E. Kingston, A. Shkumatava, I. Ulitsky, other current and former members of the Bartel lab, and A. Granger for helpful discussions; A. Liu for technical assistance; T. Eisen for hippocampal cultures; S. Eichhorn, J. Kwasniewski, K. Lin, J. Morgan, and X. Wu for scripts and computational advice. This research was supported by the NIH grants GM061835 and GM118135 (D.P.B), and T32CA009216 (B.K.). D.P.B is an investigator of the Howard Hughes Medical Institute.

---

**Figure S1.**
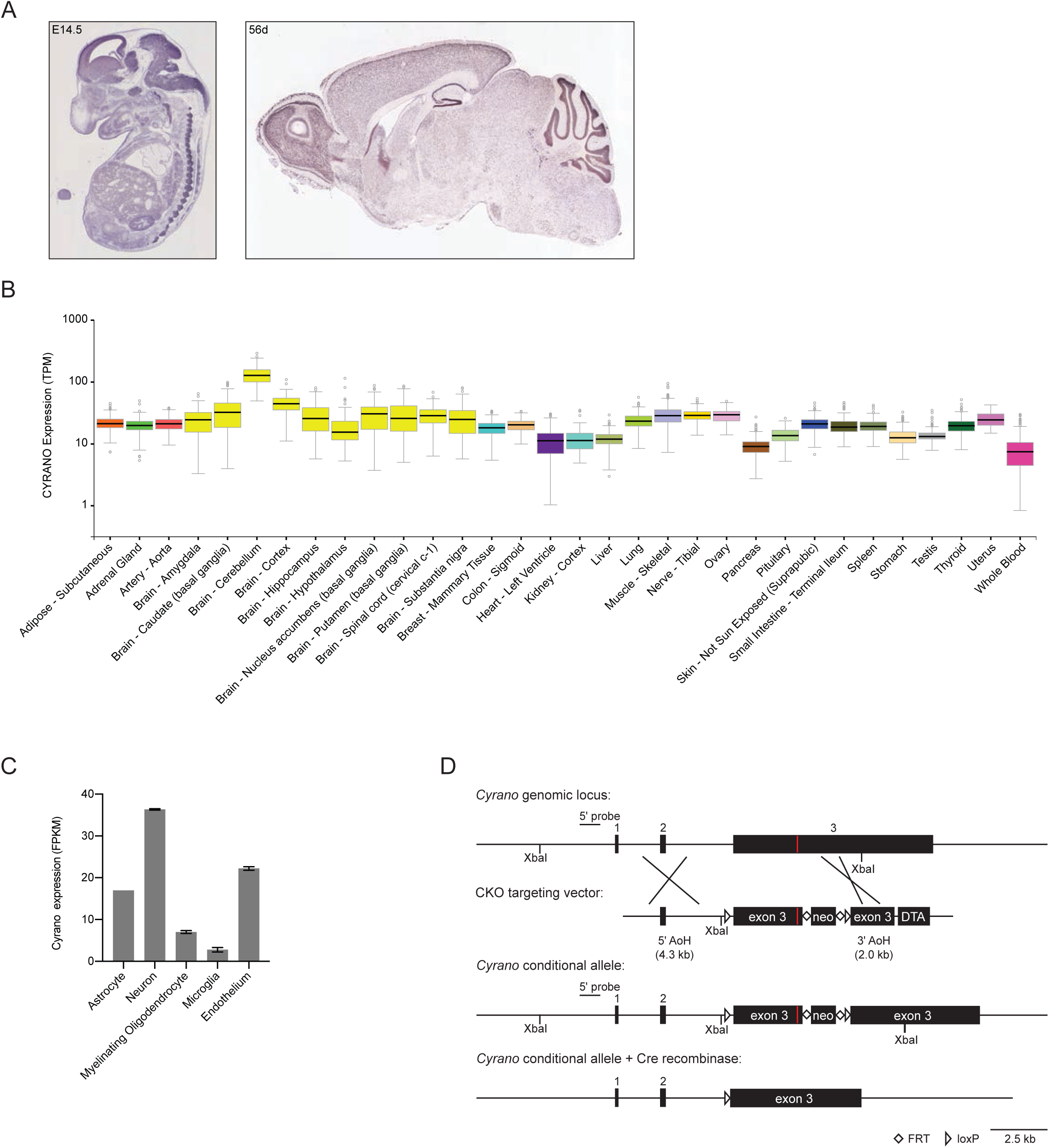
Characterization of Cyrano Expression, Related to Figure 1. (A) In situ hybridizations for Cyrano in the developing mouse embryo (E14.5) and adult mouse brain (P56) derived from GenePaint.org (Visel et al., 2004) and the Allen Brain Atlas (Lein et al., 2007). In both databases, *Cyrano* is annotated as the RIKEN cDNA *1700020I14Rik* gene. Cyrano expression from RNA-seq of 32 post-mortem human tissues (GTEx Portal 01/15/2018). Expression values in TPM (transcripts per million) are shown as box plots (box, 25^th^, median, and 75^th^ percentiles; whiskers, 1.5 times the interquartile range). In the GTEx Portal, *Cyrano* is annotated as *OIP5-AS1*. Cyrano expression from RNA-seq of purified cell populations from the mouse cortex (Dong et al., 2015). Mean expression values in FPKM (fragments per kb per million reads) are shown (error bars, range; n = 2 biological replicates for each cell type). Cyrano gene targeting strategy. Shown between the wild-type locus and the recombinant alleles is the vector used to target the first half of exon 3 (black boxes, exons; red boxes, miR-7 binding site; open triangles, loxP sites; open diamonds, FRT sites; neo, neomycin-resistance cassette; DTA, Diphtheria Toxin A; AoH, Arm of Homology). Screening of ES cell clones was performed by long-range PCR as well as Southern blot of XbaI-digested genomic DNA with a 5′ probe upstream of exon 1.

**Figure S2.**
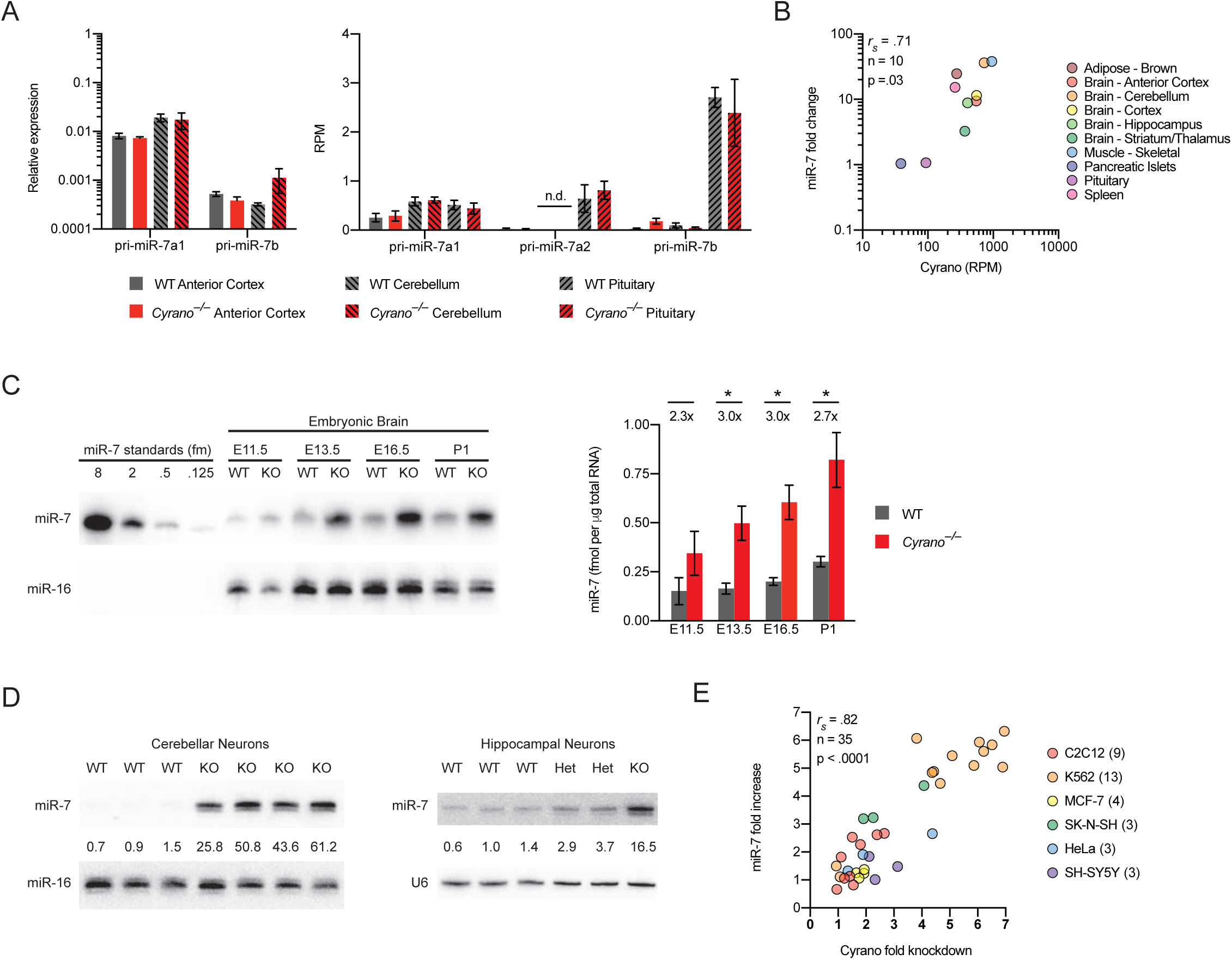
Cyrano Promotes miR-7 Destruction, Related to Figure 2. (A) The influence of Cyrano on pri-miR-7 expression in the brain. The plot on the left shows mean pri-miR-7a1 and pri-miR-7b levels in wild-type (gray bars) and *Cyrano*^*–/–*^ (red bars) anterior cortex and cerebellum (key), as determined by RT-qPCR, normalized to the geometric mean of the values for four housekeeping genes (error bars, standard deviation; n = 3 biological replicates per genotype for each tissue). The plot on the right shows mean expression values in RPM (reads per million) for all three pri-miR-7 transcripts in wild-type and *Cyrano*^*–/–*^ anterior cortex, cerebellum, and pituitary, as determined by RNA-seq (error bars, standard deviation; n = 4–5 biological replicates per genotype for each tissue; n.d., not detected). (B) Relationship between miR-7 changes observed after disrupting Cyrano and the level of wild-type Cyrano expression. Shown are miR-7 fold changes from Figure 2C plotted as a function of Cyrano expression in wild-type tissues, as determined from RNA-seq data also used for the analyses of Figure 5C. (C) The influence of Cyrano on miR-7 expression in developing mouse brain. Shown at the left is a representative RNA blot measuring miR-7 levels in wild-type and *Cyrano*^*–/–*^ brain at the indicated developmental time points. Plotted on the right are miR-7 levels in wild-type (gray bars) and *Cyrano*^*–/–*^ (red bars) embryonic brain normalized to miR-16 (error bars, standard deviation; n = 3 biological replicates per genotype). Statistically significant fold changes are indicated (*, p < .05, unpaired two-tailed t-test). (D) The influence of Cyrano on miR-7 expression in cultured primary neurons. Shown are RNA blots for wild-type and *Cyrano*^*–/–*^ DIV11 cerebellar neurons (left) and DIV12 hippocampal neurons (right). Levels of miR-7 (upper panels) were normalized to those of either miR-16 (lower panel, left) or U6 (lower panel, right), and reported relative to the mean wild-type level. (E) The relationship between the magnitude of miR-7 fold change and extent of CYRANO knockdown. Shown are miR-7 fold changes, measured by RNA blot, plotted as a function of CYRANO knockdown, as determined by RT-qPCR, for 35 mammalian CRISPRi cell lines stably expressing dCas9-KRAB and an sgRNA targeting *CYRANO*. Parental cell lines are indicated (key). For each cell line receiving a targeting sgRNA, the levels of CYRANO and miR-7 were normalized to levels of five housekeeping genes and miR-16, respectively, and shown relative to the non-targeting control.

**Figure S3.**
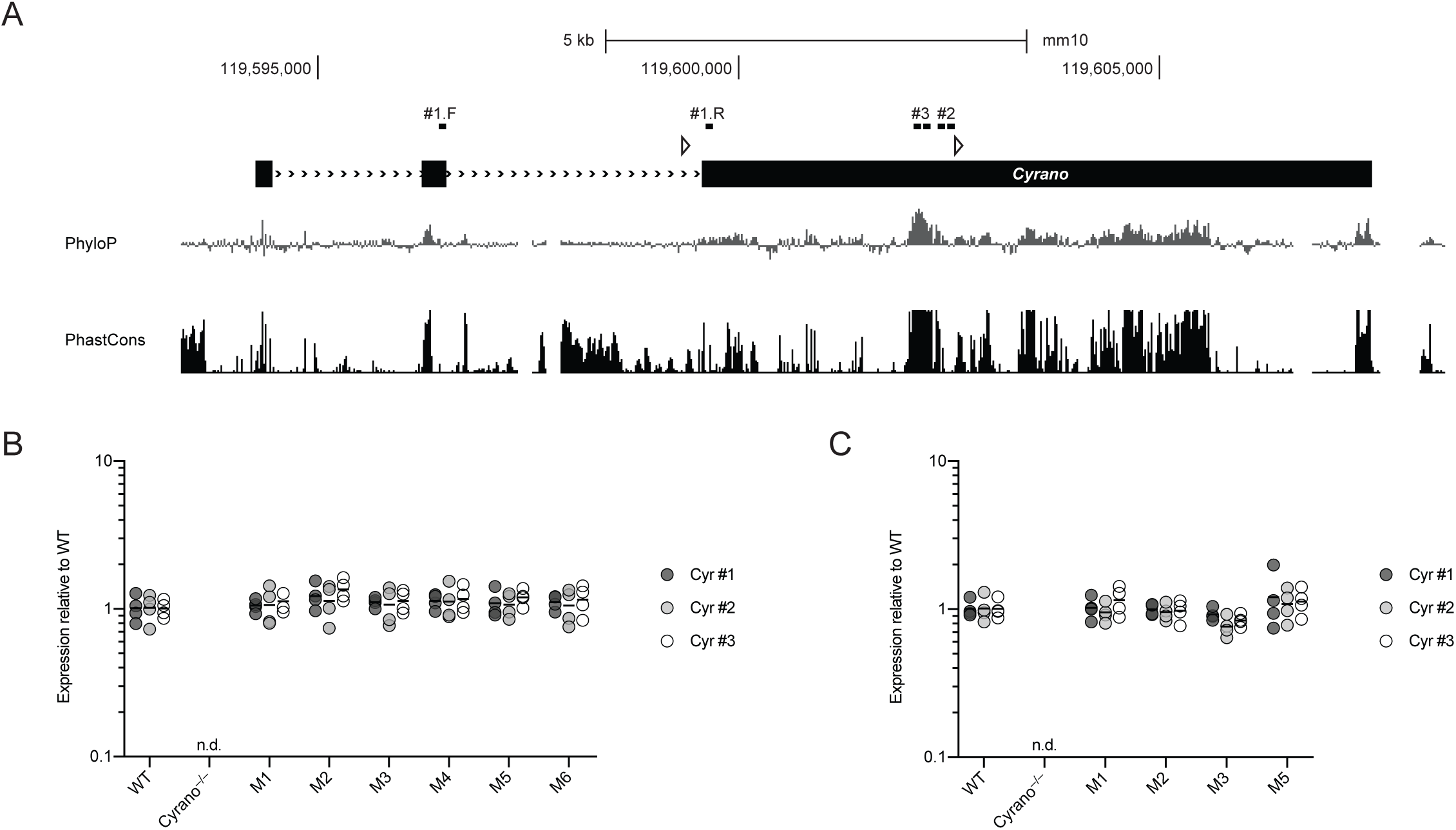
The miR-7 Site in Cyrano Does Not Destabilize Cyrano, Related to Figure 3. (A) *Cyrano* locus annotated with binding sites of 3 RT-qPCR primer pairs (#1, #2, #3). Otherwise, this panel is as in Figure 1C. (B) Cyrano expression in wild-type and *Cyrano*-mutant cerebellum. Plotted are relative Cyrano levels in wild-type and *Cyrano*-mutant cerebellum, as determined by RT-qPCR with three primer sets pairing to sites mapped in (A). Cyrano levels, normalized to that of *Actb*, are shown as fold changes relative to the mean wild-type expression (n = 4 biological replicates; n.d., not detected). (C) Cyrano expression in wild-type and *Cyrano*-mutant pituitary. Otherwise, this panel is as in (B).

**Figure S4.**
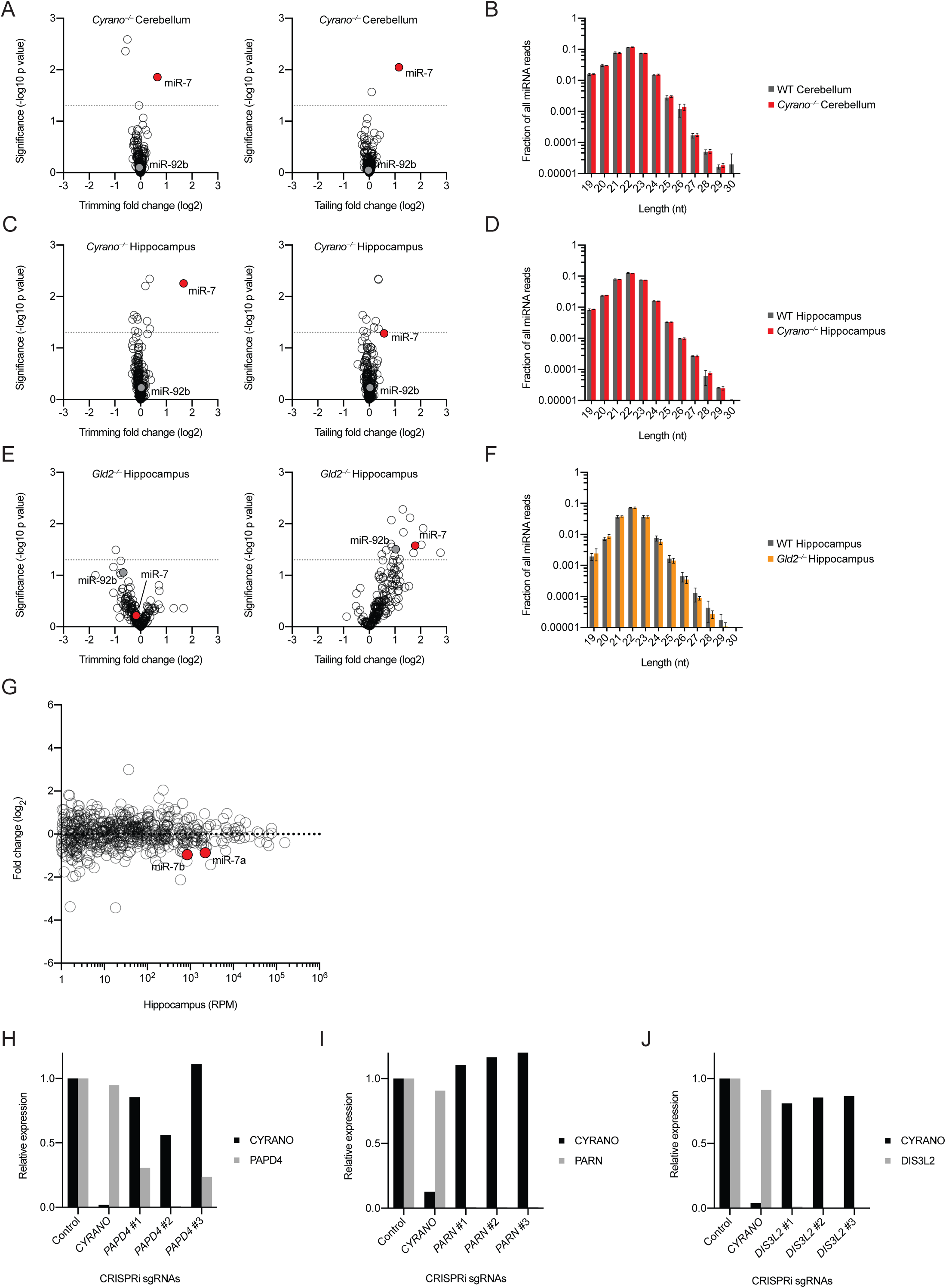
Tailing Can Be Uncoupled from Trimming and Degradation, Related to Figure 4 (A–B) The influence of Cyrano on global miRNA tailing and trimming in the cerebellum. Shown in (A) are aggregate trimming (left) and tailing (right) log_2_-fold changes for each expressed miRNA after disrupting Cyrano, plotting statistical significance of the change (based on p values determined by unpaired two-tailed t-tests, one per miRNA) as a function of the change in WT divided by *Cyrano*^*–/–*^. Each circle represents one miRNA, showing results for all miRNAs expressed above 100 RPM (read per million miRNA reads) in wild-type cerebellum. Circles representing miR-7 (miR-7a and miR-7b were combined for this analysis) and miR-92b are colored red and gray, respectively. Dotted lines indicate a p value of .05. Plotted in (B) are the length distributions of reads for all miRNAs expressed above 100 RPM (excluding miR-7) in wild-type (gray bars) and *Cyrano*^*–/–*^ (red bars) cerebellum. (C–D) The influence of Cyrano on global miRNA tailing and trimming in the hippocampus. Otherwise, these panel are as in (A–B). (E–F) The influence of Gld2 on global miRNA tailing and trimming in the hippocampus, evaluated using the data from Mansur et al. (2016). Otherwise, these panels are as in (A–B).The influence of Gld2 on miRNA levels in the hippocampus. Shown are log_2_-fold changes in mean miRNA levels observed between wild-type and *Gld2^−/–^* hippocampus, as determined by small-RNA-seq (Mansur et al., 2016) and plotted as a function of expression in wild-type hippocampus (n = 6 biological replicates per genotype). Each circle represents one miRNA, showing results for all miRNAs expressed above 1 RPM (read per million miRNA reads) in wild-type hippocampus. Circles representing miR-7 paralogs are colored red. (G) Reduced *PAPD4* expression in stable *PAPD4*-knockdown K562 cells. Shown are *CYRANO* and *PAPD4* levels, as determined by RT-qPCR, for cells expressing control sgRNA or sgRNAs targeting *CYRANO* or *PAPD4*. CYRANO and *PAPD4* levels were normalized to the geometric mean of the values for five housekeeping genes and plotted relative to levels in the control cell line. The level of *PAPD4* in line #2 was < 2% of that in the control. (H) Reduced *PARN* expression in stable *PARN*-knockdown K562 cells. The levels of *PARN* in lines #1–3 were each < 1% of that in the control. Otherwise, this panel is as in (H). (I) Reduced *DIS3L2* expression in stable *DIS3L2*-knockdown K562 cells. The levels of *DIS3L2* in lines #2–3 were < 1% of that in the control. Otherwise, this panel is as in (H).

**Figure S5.**
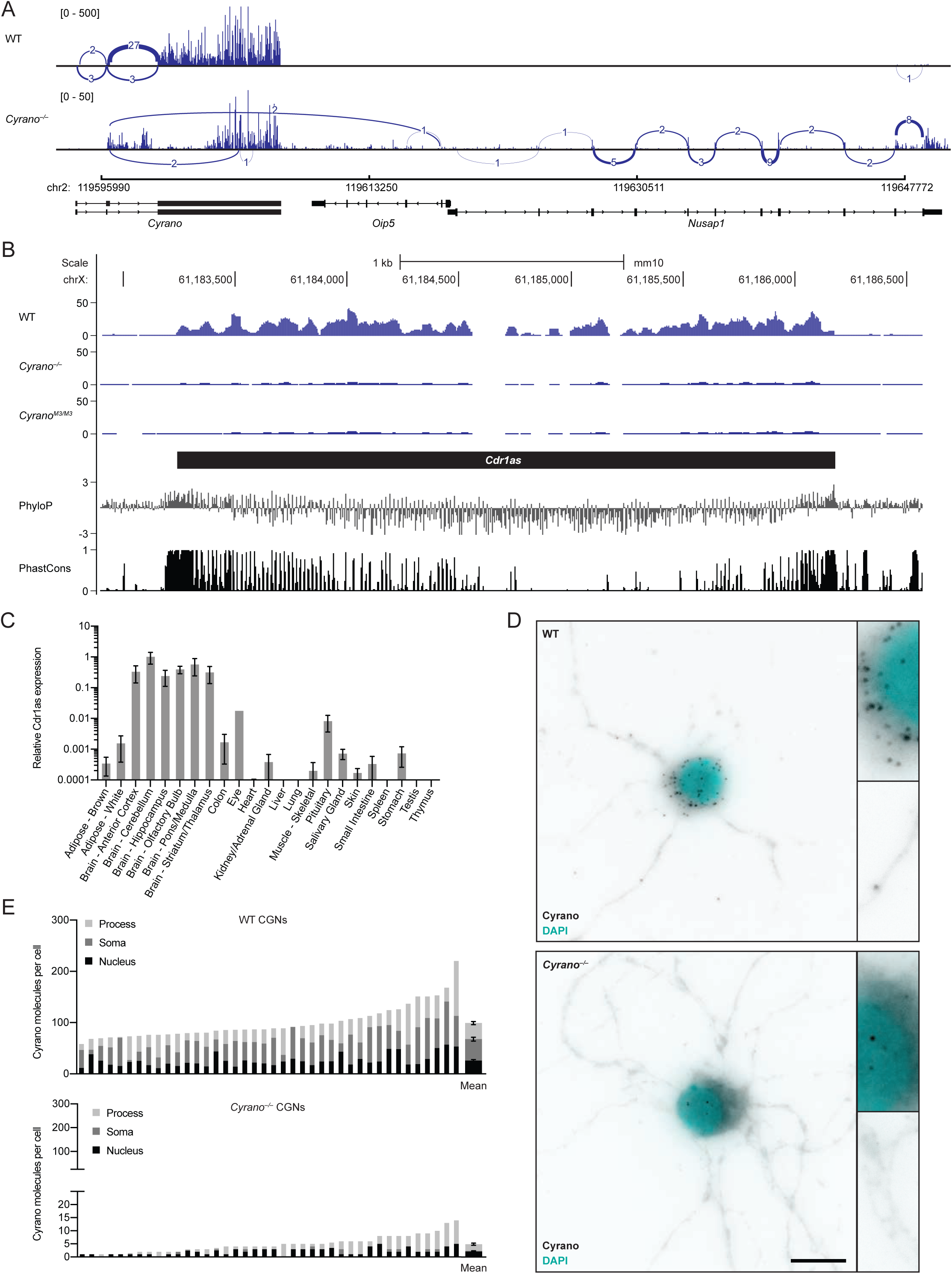
Cyrano Prevents Both the Repression of miR-7 Target mRNAs and the Destruction of Cdr1as, Related to Figure 5. (A) Sashimi plot mapping splice junctions of Cyrano and *Nusap1* transcripts in wild-type and*Cyrano*^*–/–*^ cerebellum. (B) Organization, conservation and expression of the murine Cdr1as locus. Shown below the gene model (black box, exon), are conservation plots (PhyloP and PhastCons), which are based on a 40-genome placental mammal alignment generated relative to the mouse locus (Pollard et al., 2010; Siepel et al., 2005). Shown above the gene model are RNA-seq tracks for wild-type, *Cyrano*^*–/–*^ and *Cyrano*^*M3/M3*^ cerebellum derived from libraries with similar sequencing depth. (C) Relative Cdr1as expression in the indicated tissues of 2-month-old C57Bl/6J mice. Shown are mean Cdr1as levels, as determined by RT-qPCR, normalized to the geometric mean of the values for four housekeeping genes and relative to cerebellum (error bars, range; n = 1 biological replicate for eye and lung; n = 2 biological replicates for the other 21 tissues). (D) Shown are representative images from single-molecule FISH experiments probing for Cyrano in wild-type (upper) and *Cyrano*^*–/–*^ (lower) CGNs, with insets showing increased magnification of portions of the soma and processes. Otherwise, this panel is as in Figure 5G. (E) Quantitation of single-molecule FISH results obtained when probing for Cyrano in DIV11 wild-type (upper) and *Cyrano*^*–/–*^ (lower) CGNs. Otherwise, this panel is as in Figure 5H.

**Figure S6.**
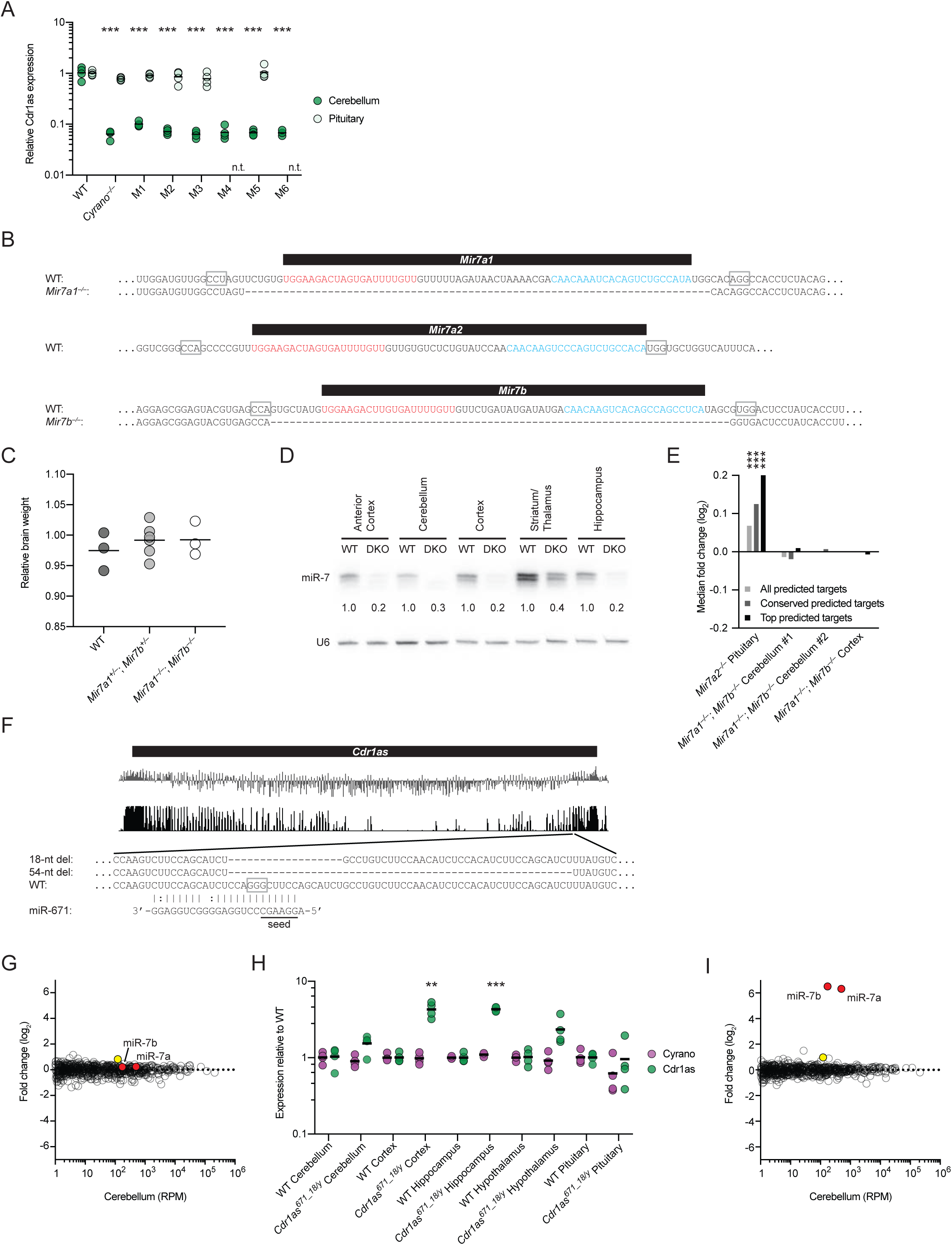
Cdr1as is Regulated by miR-7 and miR-671, Related to Figure 6. (A) The importance of the miR-7 complementary site for the ability of Cyrano to influence Cdr1as levels in cerebellum. Cdr1as expression in wild-type and Cyrano mutant cerebellum (green) was determined by RT-qPCR and plotted as in Figure 6D. For comparison, Cdr1as expression was also examined in the pituitaries (mint) of wild-type and some mutant mice (n.t., not tested). Otherwise, this panel is as in Figure 6D. (B) Schematic of the three murine Mir7 loci (black bars, pre-miRNAs; mature miRNAs and miRNA* strands are colored red and blue, respectively). The PAM motifs used for Cas9-mediated mutagenesis are outlined (open gray boxes), and nucleotides deleted in *Mir7a1* and *Mir7b* mutants are indicated by dashes. (C) Relative brain weights in P0 wild-type, *Mir7a1^+/–^*; *Mir7b^+/–^*, and *Mir7a1^−/–^*; *Mir7b^−/–^* mice from three litters produced by crosses of double heterozygotes. Weights were mean-normalized for each litter. (D) miR-7 expression in wild-type and *Mir7a1^−/–^*; *Mir7b^−/–^* (DKO) brain. Shown are RNA blots measuring miR-7 levels in the indicated brain regions of wild-type and *Mir7a1^−/–^*; *Mir7b^−/–^* mice. Levels of miR-7 (upper panel) were normalized to those of U6 (lower panel) and reported relative to the wild-type level in each region. (E) The influence of miR-7 on the predicted targets of miR-7 in the cerebellum and cortex. #1 and #2 refer to two independent experiments (n = 3–4 biological replicates per genotype for each experiment). Otherwise, this panel is as in Figure 5C. (F) Schematic of the murine Cdr1as locus. Shown below the gene model (black box, exon), are conservation plots (PhyloP and PhastCons), which are based on a 40-genome placental mammal alignment generated relative to the mouse locus (Pollard et al., 2010; Siepel et al., 2005). Pairing of miR-671 to its slicing-competent site is diagrammed below the conservation plots. The PAM motif used for Cas9-mediated mutagenesis is outlined (open gray box), and nucleotides deleted in the two different Cdr1as mutants are indicated by dashes. (G) miRNA profiling in wild-type and *Cdr1as^671^_^18/y^* cerebellum. Shown are log_2_-fold changes in mean miRNA levels observed between wild-type and *Cdr1as^671^_^18/y^* cerebellum, as determined by small-RNA-seq and plotted as a function of expression in wild-type cerebellum (n = 3 biological replicates per genotype). Each circle represents one miRNA, showing results for all miRNAs expressed above 1 RPM (read per million miRNA reads) in wild-type cerebellum. Circles representing miR-7 paralogs and miR-671 are colored red and yellow, respectively. miR-671 increased 1.8 fold (adjusted p = 5.09 × 10^−7^, as determined by DESeq2), supporting the idea that the miR-671 complementary site in Cdr1as directs some degradation of miR-671 (Piwecka et al., 2017). (H) The influence of the miR-671 site on Cdr1as accumulation in brain and pituitary. Cdr1as levels were determined by RT-qPCR and plotted as in Figure 6D (green points; black bars, mean; n = 4 biological replicates per genotype). For comparison, Cyrano levels, determined in parallel, are also plotted (purple). (I) miRNA profiling in wild-type and *Cyrano*^−/–^; *Cdr1as*^671_18/y^ cerebellum. Otherwise, this panel is as in (G).

## METHODS

### Contact for Reagent and Resource Sharing

Further information and requests for resources and reagents should be directed to and will be fulfilled by the Lead Contact, David Bartel.

### Experimental Model and Subject Details

#### Mouse Husbandry

Mice were group-housed in a 12 hr light/dark cycle (light between 07:00 and 19:00) in a temperature-controlled room (21.1 ± 1.1°C) at the Whitehead Institute for Biomedical Research with free access to water and food and maintained according to protocols approved by the Massachusetts Institute of Technology Committee on Animal Care. Euthanasia was performed by CO_2_ inhalation. The ages of mice are indicated in the figure legends or methods. Only male mice were used in this study with the exception of the RNA blot in Figure S6D.

#### Generation of *Cyrano^fl/fl^* and *Cyrano*^*–/–*^ Mice

We generated mice carrying a conditional allele of the *Cyrano* gene by homologous recombination. Genomic sequences were amplified by PCR from a BAC clone (RP23-59M9; C57BL/6J; BAC-PAC Resources, Oakland, CA). We constructed a targeting vector based on the pPGKneoF2L2DTA vector (a gift from Philippe Soriano, Addgene #13445), inserting LoxP sites in intron 2 and in a poorly conserved region of exon 3. The LoxP sites were flanked by ~4.3 kb 5′ and ~2.0 kb 3′ homology arms. The NotI-linearized targeting construct was electroporated into hybrid 129S6;C57Bl/6J ES cells. ES cell colonies were screened for correct targeting by PCR and Southern blotting, and correctly targeted ES cell clones were aggregated with albino CD-1 embryos. Chimeras were bred with FLPe transgenic mice (Cat #003946, Jackson Laboratories) to excise the neomycin resistance cassette. *Cyrano^fl/+^* mice were crossed to CMV-Cre transgenic mice (Cat #006054, Jackson Laboratories) to generate *Cyrano^+/–^* mice and then bred away from the FLPe and Cre transgenes. *Cyrano^fl/+^* and *Cyrano^+/–^* mice were each backcrossed ≥8 generations into C57Bl/6J (Cat #000664, Jackson Laboratories).

#### Generation of Transgenic Mice using Cas9

*Cyrano* miR-7 site mutant mice were generated by injecting one-cell C57Bl/6 embryos with Cas9 RNA and an sgRNA designed to cut within the miR-7 site (Figure 3A, Table S7). F0 mice containing one or more indels in the miR-7 site were crossed to wildtype C57Bl/6 mice to generate heterozygotes and then heterozygotes with the same allele were bred to homozygosity. In total, we generated 6 unique alleles, M1–M6 (Figure 3A).

*Mir7a1^−/–^*; *Mir7b^−/–^* mice were generated by injecting one-cell C57Bl/6J embryos with Cas9 RNA and 6 sgRNAs—one pair of sgRNAs flanking the pre-miRNA hairpin for each of three miR-7 loci (Figure S6b, Table S7). Of 102 F0 mice, we had one mouse with homozygous 73 and 72 bp deletions of *Mir7a1* and *Mir7b* and several mice with heterozygous deletions of *Mir7a1* and/or *Mir7b*. We did not generate any deletion alleles of *Mir7a2*. We crossed the double-knockout founder to C57Bl/6J and then began intercrossing double heterozygous F1s to generate *Mir7a1^−/–^*; *Mir7b^−/–^* mice.

*Cdr1as* miR-671 site mutant mice were generated by injecting one-cell C57Bl/6J embryos with Cas9 RNA and an sgRNA designed to cut within the miR-671 site (Figure S6b, Table S7). F0 mice containing one of two larger deletions (18 and 54 nts) in the miR-671 site were bred directly to *Cyrano^+/–^* mice (backcrossed ≥8 generations onto C57Bl/6J).

#### Genotyping

Genomic DNA was extracted from mouse earsnips using the HotSHOT method (Truett et al., 2000). For *Cyrano* miR-7 mutant mice, PCR was performed with KAPA HiFi HotStart ReadyMix (Roche) using primers indicated in Table S7 and the amplicon was purified (QiaQuick PCR Purification Kit, Qiagen) and submitted for Sanger sequencing. For all other transgenic mice, PCR was performed with KAPA 2G Fast Genotyping Mix (Roche) using primers indicated in Table S7 and the amplicon(s) were analyzed by agarose gel electrophoresis. PCR conditions and expected amplicon sizes are also indicated in Table S7.

#### Cell Lines and Cell Culture

All cells were cultured at 37°C with 5% CO_2_. Human cell lines were cultured as follows: HEK293, HEK293T, HeLa, and MCF-7 in DMEM (Gibco) with 10% FBS (Clontech); SH-SY5Y and SK-N-SH in DMEM/F12 (Gibco) with 10% FBS; K562 in RPMI-1640 (Life Technologies) with 10% FBS. Mouse cell lines were cultured as follows: C2C12 in DMEM with 10% FBS; Neuro2a in 1:1 DMEM:Opti-MEM with 10% FBS. Transfections were performed with Lipofectamine 2000 (Life Technologies) according to the manufacturer’s protocol. Each of these cell lines is of female origin.

#### CRISPRi-based Knockdown Cell Lines

Stable CRISPRi cell lines (MCF-7, SH-SY5Y, SK-N-SH, C2C12, HeLa) were generated by transduction with lentivirus carrying constitutively expressed dCas9-KRAB (Gilbert et al., 2013)(Addgene #46911). The CRISPRi K562 cell line expressed a doxycycline-inducible KRAB-dCas9 (Gilbert et al., 2014). For each experiment, a CRISPRi cell line was transduced with lentivirus expressing sgRNAs (Gilbert et al., 2014)(Addgene #60955) directed against the desired gene’s transcription start site, along with a *Cyrano*-targeting positive control sgRNA and a non-targeting negative control sgRNA. sgRNA vectors were cloned and prepared as described (Gilbert et al., 2014). sgRNA oligonucleotides are listed in Table S7. Following ~10 days of selection for sgRNA expression with puromycin (1 μg/mL in K562, 2 μg/mL in HeLa, 2.5 μg/mL in SH-SY5Y, and 3 μg/mL in MCF-7, SK-N-SH, and C2C12; Clontech), cells were harvested for RNA extraction. In the case of the inducible K562 cell line, doxycycline (Clontech) was added simultaneously with puromycin and adjusted daily to a concentration of 50 ng/mL, under the assumption of a 24-hour half-life in culture (Gilbert et al., 2014).

#### Primary Neuronal Cultures

Cerebellar granule neuron cultures were prepared from male and female P5–P6 neonates as described (Lee et al., 2009) with some modifications. Neonates were decapitated with scissors and the cerebellum and midbrain were pinched off with forceps and transferred to a petri dish containing ice-cold HBSS-glucose (Hank’s Balanced Salt Solution (Life Technologies), 0.6% glucose (w/v), 0.035% sodium bicarbonate). Under a dissecting microscope, the cerebellum was detached from the meninges and midbrain and placed in an HBSS-glucose-containing 15 mL conical tube on ice. Additional cerebella were isolated, switching petri dishes every 3 neonates, and cerebella from the same genotype were stored together.

Tissue dissociation was performed using the Papain Dissociation System kit (Worthington). Cerebella were incubated in papain solution (Earle’s Balanced Salt Solution, ~20 U/mL Papain, ~100 U/mL DNase) at 37°C for 15–20 min, gently inverting the tube every 4 min. The tissue was triturated 10–12 strokes with an FBS-coated P1000 pipet tip and the supernatant was transferred to a new tube. Cells were pelleted at 200 g for 5 min, the supernatant removed, and cells resuspended in resuspension solution (Earle’s Balanced Salt Solution, 10% albumin-ovomucoid inhibitor solution, ~100 U/mL DNase) using an FBS-coated pipet tip. The cell suspension was gently added to the top of a 15 mL conical tube containing 5 mL of albumin-ovomucoid inhibitor solution and centrifuged at 70 g for 6 min to separate intact cells from cell fragments and membranes. The supernatant was discarded and the pellet resuspended in 3–5 mL of serum-free media (Neurobasal medium, minus phenol red, supplemented with 1% GlutaMAX, 1% penicillin-streptomycin, 25 mM KCl, and 2% B-27) then passed through a 70 μM cell strainer. Cells were stained with Trypan Blue (Sigma Aldrich) and counted using a Countess (Thermo Fisher). Each cerebellum yielded 3–6 million cells with>90% viability.

Cells were plated either at ~50,000 cells/cm^2^ on #1 glass coverslips (Electron Microscopy Sciences) coated with poly-d-lysine (50 μg/mL; EMD Millipore) and laminin (2 μg/mL; Corning) and placed in 12-well plates or at ~90,000 cells/cm^2^ on tissue culture plates coated with poly-D-lysine (50 μg/mL). On DIV1, the media was exchanged for serum-free media containing 10 μM cytosine b-D-arabinofuranoside (AraC; Sigma Aldrich). On DIV3 3, the media was exchanged for serum-free media. Neurons were fed via half media exchanges every 2–3 days thereafter. Culture viability was assessed by light microscopy. A subset of cultures were also examined by DAPI staining and immunofluorescence with anti-MAP2 (1:200; EMD Millipore), anti-Tau (1:500; Synaptic Systems), and/or anti-GFAP (1:500; Dako) antibodies to ascertain the fractions of neuronal and glial cells. These cultures typically were ~90% neuronal and consisted predominantly of smaller neurons with 2–5 processes (presumably cerebellar granule neurons).

Hippocampal cultures were prepared as described (Beaudoin et al., 2012) with slight modifications. Briefly, hippocampi were dissected from P0–P1 mouse pups in ice-cold dissection media, washed twice with ice-cold dissection media, trypsinized for 20 min at 37°C, and then treated with 0.1% DNase for 2 min at room temperature. Tissue was washed again in room temperature dissection media and brought up to 10 mL in plating media. The tissue was then triturated in a 15 mL conical tube with an FBS-coated 5 mL serological pipet in 3 batches, performing no more than 10 triturations per batch and transferring the dissociated cells to a new 15 mL conical tube between each batch. Dissociated cells were passed through a 70 μM cell strainer, counted, and plated in plating media at ~100,000 cells/cm^2^ on tissue culture plates coated with poly-D-lysine (50 μg/mL). Plating media was exchanged for maintenance media between 2–6 h after plating. On DIV2, cells were treated with 5 μM AraC for 24 h and then maintenance media was fully exchanged. Neurons were fed via half media exchanges every 3 days thereafter.

### Method Details

#### RNA Extraction

For cells, total RNA was extracted with TRI Reagent (Life Technologies) according to the manufacturer’s protocol. For adherent cells, TRI Reagent was added directly to the tissue culture plate and cells were quickly detached by scraping. For suspension cells, cells were first pelleted by centrifugation. For mouse tissue, total RNA was extracted with TRI Reagent according to the manufacturer’s protocol with the following modifications. Mouse tissues were rapidly dissected after euthanasia (CO_2_) and flash frozen in Eppendorf tubes in liquid N_2_. For all but the smallest tissue (i.e. pituitary), the tissue was transferred to a 15 mL conical tube, 2 mL of TRI Reagent was added, and the tissue was homogenized with a TissueRuptor (Qiagen) and disposable probes. Phase separation was achieved with either 400 μL chloroform (J.T. Baker Analytical) or 200 μL 1-Bromo-3-chloropropane (Sigma). For samples with high fat content such as brain tissue, 2–3 volumes of 1-Bromo-3-chloropropane were necessary to fully separate the aqueous and organic layers. All RNA was resuspended in RNase-free H2O.

#### Cytoplasmic/Nuclear Fractionation

The cytoplasmic and nuclear fractions of various cell lines were isolated via mild lysis and centrifugation. Briefly, no more than 1 × 10^7^ cells were harvested and pelleted by centrifuging at 300 g for 5 min. The plasma membrane was lysed by adding 175 μL of precooled RLN+EDTA buffer (50 mM Tris pH 8.0, 140 mM NaCl,1.5 mM MgCl_2_, 0.5% NP-40, 10 mM EDTA) freshly spiked with 1 μL/mL Superasin and 1 mM DTT and gently resuspended by pipetting. Vigorous handling can result in partial nuclear lysis. Lysates were incubated on ice for 5 min then centrifuged at 500 g for 5 min to pellet nuclei. The supernatant was transferred to a new tube and spun again at 500 g for 1 min to remove any contaminating nuclei. The supernatant was transferred to a new tube containing 1 mL of TRI Reagent and vortexed vigorously. Meanwhile, the original nuclear pellet was washed once with 175 μL of precooled RLN+EDTA and spun at 500 g for 1 min. After aspirating the supernatant, the nuclei were gently resuspended in 175 μL of precooled RLN+EDTA. 1 μL was saved for microscopic confirmation of intact nuclei and the remainder was added to a new tube containing 1 mL of TRI Reagent and vortexed vigorously. The nuclear lysate was passed through a 20-gauge needle at least 5 times. 25 pg of firefly luciferase RNA (Promega) were added to each fraction.

#### Lentivirus Production and Transduction

HEK293T cells were used to package lentivirus. In 6-well tissue culture plate, ~170,000 cells/cm2 were reverse-transfected with 1.4 μg transfer plasmid, 0.94 μg pCMV-dR8.91 packaging plasmid (a gift from Jonathan Weissman), and 0.47 μg pMD2.G envelope plasmid (a gift from Didier Trono; Addgene #12259) using Lipofectamine 2000 (Thermo Fisher). After 72 h, the media was collected and centrifuged at 500 g for 10 min to pellet debris. To infect cells, 250 μL of virus-containing supernatant (~1/7^th^ of the total) was combined with 750 μL media and 8μg/mL polybrene (Santa Cruz Biotechnology) and then added to one well of a 12-well plate (containing either 200,000 suspension cells or adherent cells plated 24 h before at 25,000/cm^2^). Plates were centrifuged at 12,000 g for 90 min at room temperature then returned to the 37°C incubator.

#### RNA-seq

PolyA-selected RNA-seq libraries were prepared from 4 μg total RNA. For expression profiling of predicted miR-7 target mRNAs (Figures 5A–C), libraries were prepared using an unstranded TruSeq kit (Illumina) with tissues from 4-month-old male wild-type and *Cyrano*^*–/–*^ littermates on a mixed 129S6;C57Bl/6J background (1 sample per library; n = 4–6 samples per genotype for each tissue). For expression profiling of mRNAs and noncoding RNAs (Figure 5D, 6A, S6E), additional libraries were prepared using a stranded NEXTflex kit (Bioo Scientific) with cerebella from 1-month-old male wild-type, *Cyrano^−/–^, Cyrano^M3/M3^* and *Mir7a1^−/–^; Mir7b^−/–^* mice on a C57Bl/6J background (1 sample per library; n = 3–4 samples per genotype). All libraries were sequenced on the Illumina HiSeq platform using 40 bp single-end reads.

Reads were aligned to the mouse genome (mm10) using STAR v2.4 (Dobin et al., 2013) with the parameters “--outFilterType BySJout --outFilterIntronMotifs RemoveNoncanonicalUnannotated --outSAMtype BAM SortedByCoordinate". Aligned reads were assigned to genes using annotations from Ensembl (Mus_musculus.GRCm38.73.gtf; downloaded October 15, 2014) and htseq-count v0.6.1p1 (Anders et al., 2015) with the parameters “-m union -s no” for unstranded libraries and “-m union -s reverse” for stranded libraries. Count files were merged to generate tables of counts for each tissue organized by genotype. Differential expression was determined using DESeq2 (Love et al., 2014) with the parameter “betaPrior = FALSE". For RNA seq plots (Figure 5D, 6A), only RNAs with a mean RPM > 1 in wildtype samples and a CV < 10 standard deviations above the median CV in both wildtype and mutant samples are shown. RNA-seq browser tracks were visualized in the UCSC genome browser and IGV v2.3.34 (Robinson et al., 2011; Thorvaldsdottir et al., 2013).

The effects of miR-7 on target gene expression were determined by comparing fold changes in RNA measurements observed for genes containing miR-7 sites in their 3′ UTR to relative to those observed for control genes. The set of all predicted miR-7 targets was downloaded from TargetScanMouse v7.1 (Agarwal et al., 2015) and subsetted by conservation and by cumulative weighted context++ score to generate a set of conserved predicted targets and a set of top predicted targets, respectively. The set of controls consisted of protein-coding genes lacking a 6-mer, 7-mer, or 8-mer site to miR-7 anywhere in the transcribed region. For each tissue, only genes expressed at > 50 mean counts (approximately equivalent to 5 RPM), as determined by DESeq2, were analyzed. Genes with ≥1 miRNA site generally have longer 3′ UTRs than no site genes. To minimize fold change effects caused by 3′ UTR length differences (rather than the presence or absence of a miRNA site), we normalized the fold changes of all genes based on their 3′ UTR lengths. To do this, we first computed a linear regression from the correlation between fold changes and 3′ UTR length for no site genes. The slope of this regression was then used to normalize the fold changes for all genes.

#### Small-RNA seq

Small-RNA sequencing was performed as described (Fang and Bartel, 2015), except for size selection, which was for small RNAs from 18–32 nts (instead of 18–30 nts). A step-by-step protocol is available at http://bartellab.wi.mit.edu/protocols.html. Libraries were prepared with 5 μg total RNA from tissues of 1-month-old male mice on a C57Bl/6J background. Libraries were sequenced on the Illumina HiSeq platform using 50 bp single-end reads. Reads were trimmed of adaptor sequence using cutadapt (Martin, 2011) and filtered using fastq_quality_filter (FastX Toolkit; http://hannonlab.cshl.edu/fastx_toolkit/) with the parameters “-q 30 –p 100” to ensure that all bases have an accuracy of 99.9%.

To count the miRNAs in each library, the first 18 nts of each read were string-matched to a dictionary of miRNA sequences. Mouse and human miRNA dictionaries were based on annotations in miRbase_v20 and miRbase_v21, respectively. Matching to the first 18 nts has the advantage of combining all miRNA isoforms into a single count but also results in ambiguously assigning reads when two distinct mature miRNAs are identical for the first 18 nts but differ at one or more positions in the 3′ end (such as let-7a-5p and let-7c-5p). To avoid this ambiguity, these miRNAs (<5% of all annotations) were removed from the dictionary. Count files were merged to generate tables of counts for each tissue organized by genotype. Differential expression was determined using DESeq2 (Love et al., 2014) with the parameter “betaPrior = FALSE". For small-RNA seq plots (Figure 2A–B, 2D, S6G–H), only miRNAs with a mean RPM > 1 in wildtype samples and a CV < 5 standard deviations above the median CV in both wildtype and mutant samples are shown.

For our tailing and trimming analyses, we used the same string-matching described above while also binning reads by length from 19–30 bases. For each miRNA with a mean RPM > 100, we calculated the fraction of reads for each length by dividing by the total number of reads. Any length with a fraction > 25% was considered “mature.” Trimmed reads were all reads shorter than the shortest mature length and tailed reads were all reads longer than the longest mature length. In this fashion, we were able to determine the aggregate number of trimmed and tailed reads for each miRNA. To identify the nucleotide(s) added during tailing, we used custom scripts to count all homopolymers appended to mature miR-7 (both the 23- and 24-nt isoforms).

#### RNA Blots

1–5 μg of total RNA for each sample was resolved on a denaturing polyacrylamide gel and transferred onto a Hybond-NX membrane (GE Healthcare) using a semi-dry transfer apparatus (Bio-Rad). Because UV crosslinking is biased against shorter RNAs, EDC (*N*-(3-dimethylaminopropyl)-*N*’-ethylcarbodiimide; Thermo Scientific) was used to chemically crosslink 5′ phosphates to the membrane (Pall et al., 2007). Blots were hybridized to radio-labeled DNA or LNA probes. Probe oligonucleotides are listed in Table S7. RNA blot data were analyzed with ImageQuant TL (v8.1.0.0). A step-by-step protocol is available at http://bartellab.wi.mit.edu/protocols.html. For some experiments, synthetic miR-7 standards were run in lanes adjacent to the experimental samples to generate a standard curve.

#### RT-qPCR

For mouse tissues, 1 μg total RNA was treated with RQ1 DNase (Promega) or Turbo DNase (Life Technologies), then reverse transcribed with Superscript III (Life Technologies) and random hexamers. Expression of Cyrano and Cdr1as (see Table S7 for primer sequences) were measured by qPCR using SYBR Green (Applied Biosystems) on a QuantStudio 6 system (Applied Biosystems) and quantified using the ΔΔCT method with Actb expression or the geometric mean of Actb, U1, U2, and U6 expression as internal normalization controls.

For stable CRISPRi cell lines, 1 μg total RNA was reverse transcribed with QuantiTect RT Kit (Qiagen) followed by qPCR using SYBR Green. Expression of CYRANO and other mRNAs (see Table S7 for primer sequences) were quantified using the ΔΔCT method with the geometric mean of GAPDH, ACTB, U1, U2, and U6 expression as an internal normalization control and the non-targeted sample as the inter-sample normalization control.

#### Absolute quantification of miR-7, Cyrano, and Cdr1as in CGNs and cell lines

To quantify the number of molecules of a given RNA per cell, we first needed to determine the amount of total RNA per cell. For this, total RNA was extracted from a defined number of cells (RNA loss was assumed to be negligible). For human cell lines, cell number was determined using a Countess cell counter. For mouse DIV11 CGN, cell number was calculated from light micrographs of 6-well plates (10 20x images per sample) prior to RNA extraction. By this method, we estimated mean total RNA per cell as follows: 7.7 pg per CGN, 19.9 pg per HEK293T, and 23.2 pg per HeLa (n = 3 samples per cell type; Table S4).

Next, we generated standards for miR-7, Cyrano, and Cdr1as. For miR-7, 10 pmol of a 23-mer RNA oligo (IDT) matching the human miR-7 and mouse miR-7a sequence was PNK-labeled with cold ATP, desalted on a P30 column (Bio-Rad) and then serially diluted in 200 ng/μL yeast tRNA (Life Technologies). For Cyrano, the first 3.5 kb of the mouse Cyrano cDNA and the first 3.7 kb of the human CYRANO cDNA were PCR’d to add a 5′ T7 sequence and then gel-extracted (Qiaquick Gel Extraction, Qiagen). 1 pmol of each PCR product was in vitro transcribed with T7 MEGAscript (Ambion), according to manufacturer’s protocol, and purified with Turbo DNase and MEGAclear (Ambion). In addition to the full-length RNA, in vitro transcription reactions often contain shorter products, which could confound accurate quantification. To ensure that the standards consisted only of full-length RNA, we size-selected the RNA on a native RNase-free 1% agarose gel run with RiboRuler HR ladder (Thermo Scientific) according to the protocol of Masek et al. (2005). Size-selected RNA was purified with Zymoclean Gel RNA Recovery Kit (Zymo Research), and quantified using a NanoDrop spectrophotometer (Thermo Scientific). For Cdr1as, a 243 bp fragment of mouse Cdr1as and 362 bp fragment of human CDR1AS (both containing the back-splice junction), each prepended with a 5′ T7 sequence, were ordered from IDT as a gBlock (see Table S7 for sequences). ~1 pmol of dsDNA was in vitro transcribed with T7 MEGAshortscript (Ambion), according to manufacturer’s protocol, and then cleaned up with MEGAclear. The full-length RNAs were size-selected on a 4% polyacrylamide gel run with RNA Century markers (Ambion), eluted overnight at 4°C, purified with a Spin-X column (Corning), concentrated by EtOH precipitation, and quantified by NanoDrop. The molecule weight of each RNA standard was calculated based on its sequence (1503509.3 g/mol for mouse Cyrano; 1147517.4 g/mol for human Cyrano; 70377.1 g/mol for mouse Cdr1as; 108587.7 g/mol for human Cdr1as) and RNA standards were serially diluted, 10^10^ to 10^4^ copies, in 1 μg total RNA derived from HeLa cells (for mouse standards) or 3T3 cells (for human standards).

To determine absolute molecules of miR-7, 5 μg total RNA from CGN and HEK293T and 15 μg total RNA from HeLa were analyzed by RNA blot alongside a serial dilution of synthetic miR-7 standards. To determine absolute molecules of Cyrano and Cdr1as, RT-qPCR with Superscript III and gene-specific primers (Table S7) was performed on 1 μg total RNA from CGN, HEK293T, and HeLa alongside serial dilutions of in vitro transcribed standards.

#### Single-molecule FISH on CGNs

Single-molecule FISH was performed on cerebellar granule neurons grown on glass coverslips according to the following protocol, adapted from (Batish et al., 2011). Cells were washed twice in PBS and fixed in freshly-made 4% paraformaldehyde (Electron Microscopy Sciences) in PBS for 10 min. The fixation solution was aspirated and cells were gently washed in PBS then stored in 70% ethanol for ≥ h (up to 2 weeks) at 4°C.

Coverslips were equilibrated for ≥2 min in washing buffer (10% formamide, 2X SSC).Custom probes 3′ end-modified with Quasar 670 dye (see Table S7 for sequences) were diluted to 25 nM in hybridization solution (10% formamide, 2X SSC, 100 mg/mL dextran sulfate) and 50μL of probe/hybridization mix was dotted on parafilm pressed to a glass plate, 1 dot per coverslip. Coverslips were carefully removed from washing buffer with forceps and then placed on the probe/hybridization mix with the cell side facing the liquid. The glass plate was incubated in a humidifying chamber, protected from light, at 37°C overnight. Subsequently, the coverslips were washed 30 min in washing buffer to remove excess probe, 30 min in washing buffer containing 100 ng/mL DAPI, and 5 min in washing buffer to remove excess DAPI, always protected from light. Coverslips were mounted on glass slides with ProLong Gold (Invitrogen) and allowed to cure overnight, protected from light, before sealing with clear nail polish.

Images were acquired using a 100x oil-immersion objective with a numerical aperture of 1.4 on a wide-field fluorescence microscope (either a Zeiss AxioPlan2 with cooled CCD camera or a GE Healthcare Deltavision with sCMOS camera). In each field, 10–30 z-sections with 0.2 μm spacing were acquired. On the AxioPlan2, exposure times were 2 s Cy5, 0.5 s GFP, 0.5 s DAPI. On the Deltavision, exposure times were 0.5 s Cy5 (100% intensity), 0.3 s GFP (100%),0.01 s DAPI (50%).

For quantification, hot pixels were removed from individual stacks using Remove Outliers (radius 1 pixel, threshold 10) and Despeckle functions in ImageJ. Paired Cy5 and GFP stacks were loaded into the MATLAB program ImageM (Lyubimova et al., 2013), z-projected, and number of dots in each z-projection was enumerated using the Count Dots feature. Background dots, defined as dots counted in both the Cy5 and GFP stacks, were subtracted. The number of dots was divided by the number of intact, non-pyknotic nuclei in each image to generate average molecules per cell for each field. In total, 40 fields were analyzed for each probe and each genotype and plotted in Figure 5F. To quantify the number of dots by subcellular location, the Segmentation feature in ImageM was used to outline either the soma using GFP autofluorescence or the nucleus using DAPI. In this fashion, the number of dots in the nucleus was determined directly, the number of dots in the cytoplasmic soma—here, referred to as soma in Figures 5H and S4D—was calculated by subtracting nuclear dots from soma dots, and the number of dots in neuronal processes was calculated by subtracting soma dots from total dots per field. It is also worth noting that because quantification is performed on z-projected images rather than individual sections in a z-stack, the number of dots assigned to the nucleus is likely an overestimation.

For the generation of publication quality images, hot pixels were removed as described above, stacks were z-projected using Max Intensity and then combined using Merge function in ImageJ. Merged files were converted to an RGB file and imported into Photoshop (Adobe) for cropping.

### Quantification and Statistical Analysis

Graphs were generated in GraphPad Prism 7. Statistical parameters including the value of n, statistical test, and statistical significance (p value) are reported in the Figures and the Figure Legends. Biological replicates refer to samples derived from different mice. No statistical methods were used to predetermine sample size.

### Data and Software Availability

Sequencing datasets generated in this study are being deposited in NCBI GEO and will be available upon publication.

**Table S1:**

|  | *Cyrano* 2A9<sup>+/+</sup> | *Cyrano* 2A9<sup>+/-</sup> | *Cyrano* 2A9<sup>-/-</sup> |
| --- | --- | --- | --- |
| **P21** | 81 | 154 | 76 |
| % observed | 26% | 50% | 24% |
| % expected | 25% | 50% | 25% |
| $\chi^2$ p=.91 | | | |

**Table S2:**

|  | *Cyrano* 1F1<sup>+/+</sup> | *Cyrano* 1F1<sup>+/-</sup> | *Cyrano* 1F1<sup>-/-</sup> |
| --- | --- | --- | --- |
| **P21** | 24 | 55 | 26 |
| % observed | 23% | 52% | 25% |
| % expected | 25% | 50% | 25% |
| $\chi^2$ p=.85 | | | |

**Table S3:**

| Cell type | pg total RNA/cell | Cyrano copies/cell | Cdr1as copies/cell | miR-7 copies/cell |
| --- | --- | --- | --- | --- |
| **CGN (DIV11)** | 7.7 ± 0.9 | 102 ± 16 | 146 ± 37 | 40 ± 7 |
| **HEK293T** | 19.9 ± 1.5 | 7 ± 2 | 6 ± 1 | 845 ± 204 |
| **HeLa** | 23.2 ± 3.7 | 22 ± 3 | Not detected | 148 ± 66 |

**Table S4:**

| *Mir7a1*: | *Mir7a1*<sup>+/+</sup> |  |  | *Mir7a1*<sup>+/-</sup> |  |  | *Mir7a1*<sup>-/-</sup> |  |  |
| --- | --- | --- | --- | --- | --- | --- | --- | --- | --- |
| *Mir7b*: | *Mir7b*<sup>+/+</sup> | *Mir7b*<sup>+/-</sup> | *Mir7b*<sup>-/-</sup> | *Mir7b*<sup>+/+</sup> | *Mir7b*<sup>+/-</sup> | *Mir7b*<sup>-/-</sup> | *Mir7b*<sup>+/+</sup> | *Mir7b*<sup>+/-</sup> | *Mir7b*<sup>-/-</sup> |
| **P21** | 9 | 13 | 8 | 17 | 33 | 19 | 5 | 18 | 8 |
| % observed | 7% | 10% | 6% | 13% | 25% | 15% | 4% | 14% | 6% |
| % expected | 6% | 13% | 6% | 13% | 25% | 13% | 6% | 13% | 6% |
| $\chi^2$ p=.95 | | | | | | | | | |

**Table S5:**
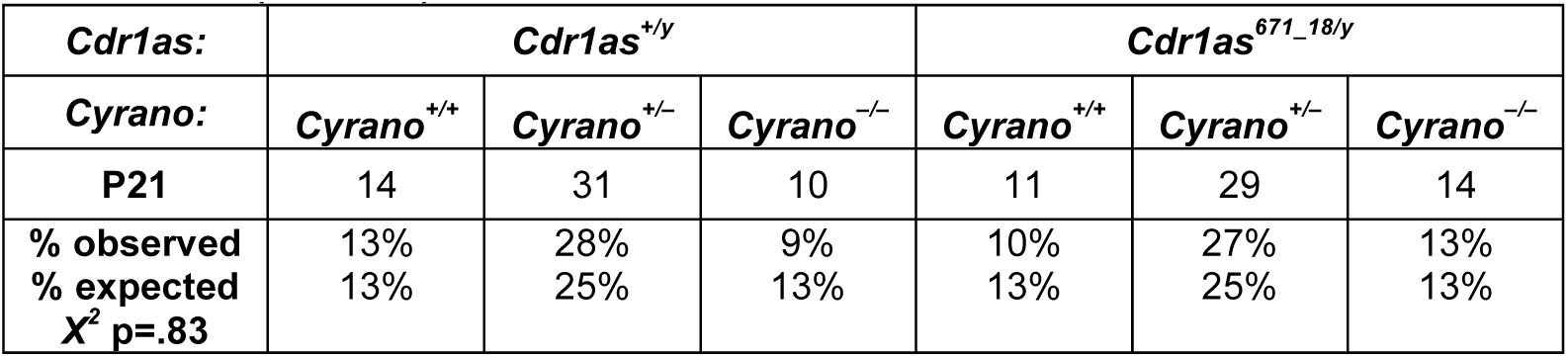
(Males)

**Table S5:** (Females)

| *Cdr1as*: | *Cdr1as*<sup>+/671_18</sup> |  |  | *Cdr1as*<sup>671_18/671_18</sup> |  |  |
| --- | --- | --- | --- | --- | --- | --- |
| *Cyrano*: | *Cyrano*<sup>+/+</sup> | *Cyrano*<sup>+/-</sup> | *Cyrano*<sup>-/-</sup> | *Cyrano*<sup>+/+</sup> | *Cyrano*<sup>+/-</sup> | *Cyrano*<sup>-/-</sup> |
| **P21** | 17 | 24 | 15 | 15 | 23 | 9 |
| % observed | 17% | 23% | 15% | 15% | 22% | 9% |
| % expected | 13% | 25% | 13% | 13% | 25% | 13% |
| $\chi^2$ p=.61 | | | | | | |

**Table S6:**
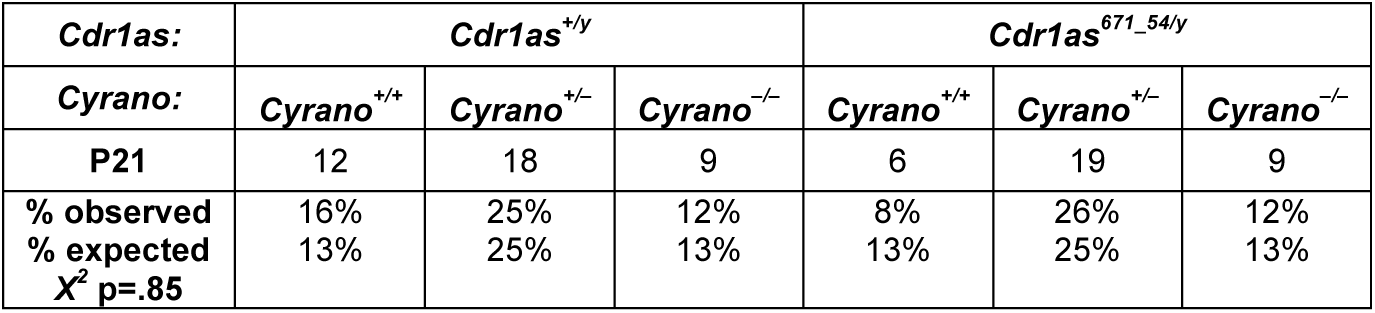
(Males)

**Table S6:**
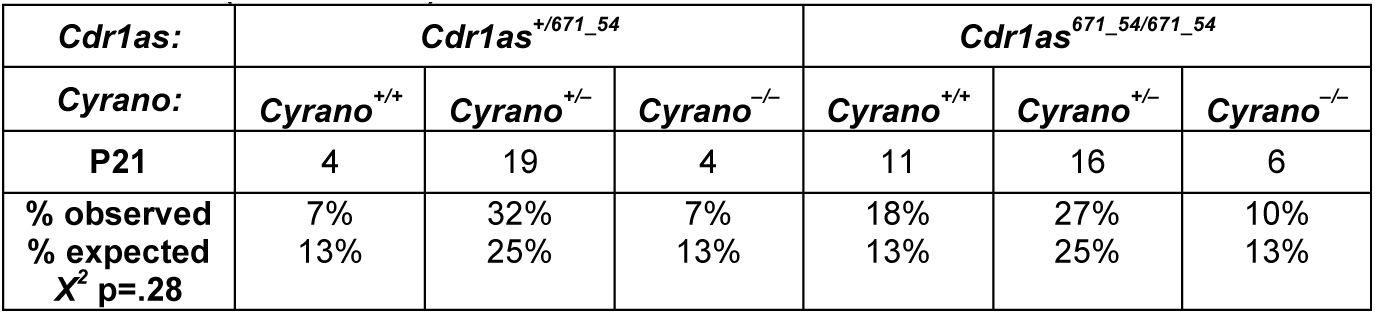
(Females)

